# Expanded gut microbial genomes from Chinese populations reveal population-specific genomic features related to human physiological traits

**DOI:** 10.1101/2024.10.19.618999

**Authors:** Quanbin Dong, Beining Ma, Xiaofeng Zhou, Pan Huang, Mengke Gao, Shuai Yang, Yuwen Jiao, Yan Zhou, Zhun Shi, Qiufeng Deng, Dongxu Hua, Xiuchao Wang, Lu Liu, Chengcheng Zhang, Chuan Zhang, Mengmeng Kong, Chang He, Tingting Wu, Huayiyang Zou, Jing Shi, Yanhui Sheng, Yifeng Wang, GMR Consortium, Liming Tang, Shixian Hu, Huanzi Zhong, Wei Sun, Wei Chen, Qixiao Zhai, Xiangqing Kong, Yan Zheng, Lianmin Chen

## Abstract

**Background:** A comprehensive and representative reference database is crucial for accurate taxonomic and functional profiling of the human gut microbiome in population-level studies. However, with 70% of current microbial reference data derived from Europeans and the Americans, East Asia, especially China, remain underrepresented.

**Methods:** We constructed the human Gut Microbiome Reference (GMR), comprising 478,588 high-quality microbial genomes from Chinese (247,134) and non-Chinese (231,454) populations. Species-level clustering and protein annotations were performed to characterize microbial diversity and function. We further integrated novel genomes into taxonomic profiling database and validated the improvements using independent cohort data.

**Results:** The GMR dataset spans 6,664 species, including 26.4% newly classified species, and encodes over 20 million unique proteins, with 47% lacking known functional annotations. Notably, we observed that 35.35% and 32.46% of species unique to Chinese and non-Chinese populations, respectively. For 2,145 species shared between populations, 74% of 304 species with balanced prevalence between populations exhibited population-specific phylogenetic stratification, involving health relevant functionalities such as antibiotic resistance. Integration of novel genomes into taxonomic improved population-level species profiling by up to 23% and uncovered replicable associations between novel species and host physiological traits.

**Conclusions:** Our study largely expands the compositional and functional landscape of the human gut microbiome, providing a crucial resource for studying the role of gut microbiome for regional health disparities.

## Background

The human gut is colonized by trillions of microbes with diverse metabolic activities that are essential for human health, and its composition is influenced by a variety of factors, such as age, diet and geographical environment[1, 2]. To precisely decipher the taxonomic and functional diversity of the human gut microbiome ecosystem, the construction of comprehensive reference genome and gene catalogs is of paramount importance. In recent years, initiatives such as the Unified Human Gastrointestinal Genomes (UHGG) consortium[3] have made significant strides by integrating multiple international sequencing efforts[4–9] to create a unified, non-redundant reference genome dataset for the human gut microbiome.

However, despite their utility, existing catalogs exhibit pronounced geographic biases. Over 70% of UHGG genomes originate from European and American populations, whereas Asian cohorts, particularly those from China, contribute less than 30% of the genomic data. This underrepresentation may introduce significant bias in studies focusing on Chinese cohorts, potentially leading to systematic omission of microbial taxa unique to the Chinese population due to the lack of reference sequences. Besides, the current reference resources may not fully capture the global diversity and functional heterogeneity of gut microbes. In particular, within-species genetic diversity plays a critical role in microbial metabolism, colonization, and pathogenicity[10–12]. For instance, only specific strains of *Escherichia coli* produce carcinogens[13], while others are non-pathogenic[14]. Similarly, well-known microbes like *Prevotella copri*[15–18] and *Akkermansia muciniphila* [19] have been shown to possess substantial phylogenetic and functional diversity across global populations. Despite these findings, systematic comparisons of strain-level genomic diversity among prominent microbes remain scarce, and the potential physiological relationships between host– or region-specific microbial strains and their functional roles are still poorly understood.

Although previous research[3, 20, 21] has made substantial efforts to expand microbial reference genome collections for the Chinese population, the current sample size remains inadequate to fully capture the diversity and complexity of the gut microbiome in China’s vast population of approximately 1.4 billion people. The resulting gap not only hampers the ability to perform robust comparative analyses between Chinese and non-Chinese populations but also limits insights into key functional aspects such as antibiotic resistance, which has profound implications for public health.

In response to these challenges, we introduces a comprehensive human Gut Microbial Reference (GMR) collection that significantly broadens the scope of current microbial resources. By integrating nearly 500,000 high-quality genomes derived from existing datasets[3, 20–23] with an additional 6,657 metagenomic samples from large Chinese cohorts, we have developed an expanded database that offers a more balanced representation of global gut microbiome diversity. This enriched collection lays the foundation for robust comparative analyses aimed at elucidating both taxonomic differences and functional gene profiles between Chinese and non-Chinese populations. Our analyses focus on examining species distribution patterns and the prevalence of functional genes, with a particular emphasis on those associated with antibiotic resistance—a critical area given the rising concern over antimicrobial resistance worldwide. Furthermore, for potential novel microbial species, we incorporated their genomes into the MetaPhlAn4[24] taxonomic classification database (released in March 2024) to facilitate population-level profiling and to delineate their compositional and genomic relationships with diverse host physiological traits. The performance and utility of this updated database have been rigorously validated using independent cohort datasets, thereby affirming its applicability in diverse microbiome research contexts.

## Methods

### Overview of the metagenomic datasets

We compiled publicly available prokaryotic genomes sampled from the Chinese gut as of February 2024. The isolates from Chinese individuals are collected from CGR2[23] (https://db.cngb.org/codeplot/datasets/CGR2) and hGMB[22] (https://hgmb.nmdc.cn/) collections, which are the largest microbial culture repositories in China. A total of 104 and 1,804 isolates were collected from these datasets, respectively. The metagenomic data were primarily obtained from studies by Huang et al.[20] (https://db.cngb.org/search/project/CNP0004122/), Qi et al.[25] (https://ehp.niehs.nih.gov/doi/10.1289/isee.2024.0796), Zhang et al.[26] (https://www.ebi.ac.uk/ena/browser/view/PRJEB65297), and Li et al.[27] (https://www.ncbi.nlm.nih.gov/sra?linkname=bioproject_sra_all&from_uid=772518). In total, we downloaded a total of 76.8 TB of data from 6,657 metagenomic samples and assembled MAGs using the methods described below. In addition, we included a recently published dataset of 6,729 MAGs from Inner Mongolians[21] (https://doi.org/10.6084/m9.figshare.19661523) and the complete set of 289,232 genomes from the UHGG[3] dataset (human gut, v2.0.1, https://ftp.ebi.ac.uk/pub/databases/metagenomics/mgnify_genomes/human-gut/v2.0.1/).

### Metagenomic *de novo* assembly and binning

To maximize the quantity of reconstructed genomes, we employed a single-sample assembly strategy following the recommendations of Stephen *et al.*[6] and Pasolli *et al.*[8]. The raw metagenomic reads were pre-processed through a pipeline with the following key steps: 1). trimming of low-quality bases and adapter sequences with Trimmomatic (v.0.33)[28]; 2). removing duplicate reads with Trf (v.4.09)[29]; 3). filtering out human-derived reads by aligning against the human reference genome GRCh37 (hg19) with KneadData (v.0.10.0)[30] integrated Bowtie2 (v.2.1.0)[31]. The clean paired-end reads were then assembled with MEGAHIT (v.1.2.9)[32]. Contigs longer than 1,000 bp were used for genome binning with MetaBAT2 (v.2.12.1)[33]. Briefly, raw reads were mapped back to the filtered contigs using Bowtie2 (v.2.1.0)[31], and coverage was calculated using the *jgi_summarize_bam_contig_depths* script. Genome binning was subsequently performed with ‘*metabat*’ script. A total of 500,154 genome bins were reconstructed from 6,657 metagenomic samples.

### Quality control of genome bins

For newly reconstructed genome bins, genome quality (completeness and contamination) was assessed using CheckM (v.1.1.3)[34] with the ‘*lineage_wf* ‘ workflow. We selected only those genomes that met the UHGG criteria: > 50% genome completeness, < 5% contamination, and an estimated quality score (completeness – 5 × contamination) > 50. Including UHGG, there are 478,588 genomes that meet these criteria and were kept for downstream analysis and included in the GMR.

### Cluster metagenomic assemblies into species-level genome bins

To address the computational burden of clustering all 478,588 genomes, we employed a two-step approach. First, we used GTDB (R214)[35, 36] to annotate all genomes at the species level. In this process, 1,731, 529, 57, 61, 69, 31, and 0 genomes from CGMR, Zheng, IMGG, PRJEB65297, PRJNA772518, CGR2, and hGMB, could not be annotated to the species level respectively. As UHGG annotations were based on the R202 version, 451 and 2,525 genomes from Chinese and non-Chinese samples, respectively, could not be assigned a species-level annotation. We re-annotated these genomes using the newer R214 version. However, 101 and 899 genomes from Chinese and non-Chinese samples, respectively, still lacked species-level annotations. Finally, we employed dRep[37] to cluster the remaining 3,478 unannotated genomes at a 95% ANI threshold with the following options: ‘*-pa 0.9*’ (primary cluster at 90%), ‘*-sa 0.95*’ (secondary cluster at 95%), ‘*-cm larger*’ (coverage method: larger), ‘*-comp 50*’ (completeness threshold of 50%), ‘*-con 5*’ (contamination threshold of 5%) and ‘*-nc 0.30*’ (coverage threshold of 0.3), resulting in 1,762 distinct clusters. Combining these clusters with the 4,902 genomes that received species-level annotations through re-annotation, we obtained a final collection of 6,664 clusters.

The best quality genome from each species cluster was selected as its representative on the basis of genome completeness, minimal contamination and assembly N50 based on the following formula[3]:

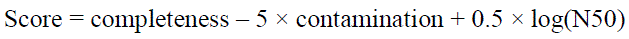

where N50 is the assembly contiguity characterized by the minimum contig size in which half of the total genome sequence is contained. The genome with the highest score was chosen as the species representative, and the final 6,664 genomes was used to generate the GMR representative catalog. The phylogenetic tree of 6,624 bacteria species was visualized and annotated with Interactive Tree Of Life (iTOL)[38] (https://itol.embl.de/itol.cgi)

### Functional annotation of GMR genomes

For functional annotation, protein-coding sequences (CDS) were predicted from each of the 478,588 genomes using Prodigal (v2.6.3)[39]. The following options were applied: ‘*-c*’ to ensure only closed-ended protein predictions, ‘*-m*’ to prevent genes from being built across stretches of sequence marked as Ns, and ‘*-p single*’ to indicate single mode for genome assemblies containing a single species.

Functional annotation of archaeal and bacterial genomes was initially performed using Prokka (v. 1.14.6)[40], with the parameters ‘*--kingdom Archaea*’ and ‘*--kingdom Bacteria*’ applied accordingly.

The nonredundant GMRP catalog was constructed from a combined set of 1,028,234,783 genes predicted across all 478,588 genomes. Protein clustering of the GMRP was performed with the ‘*cluster*’ function of MMseqs2[41] (v.7.4e23d) with options ‘*--cov-mode 1 –c 0.8* (minimum coverage threshold of 80% the length of the shortest sequence)’. The ‘*--min-seq-id* ‘ option was set at 1, 0.9 and 0.5 to generate the catalogs at 100%, 90% and 50% protein identity, respectively. The single representative nucleotide sequence from each protein clusters, obtained using the ‘*createtsv*’ function of MMseqs2, was then functionally annotated with eggNOG-mapper (v.2.1.12) using database v5.0.2[42, 43]. The COG[44] and KEGG[45] module were derived from the eggNOG-mapper results.

### Antimicrobial resistance genes

Predicted genes were annotated with The ResFinder database[46] by using abricate (https://github.com/tseemann/abricate) (v.0.5) with default parameters. We filtered the results to retain genes with coverage > 50% and identity > 80%.

### Pangenome generation

Pan-genome analyses were carried out using Roary (v3.12.0)[47]. Roary utilizes a MCL (Markov Clustering) algorithm to cluster orthologous genes based on BLASTP results and identify core and accessory genes within the genomes of interest. For all genomes in each species cluster, we constructed a pan-genome with a minimum amino acid identity for a positive match at 90% (*’-i 90’*), a core gene defined at 90% presence (*’-cd 90’*) and no paralog splitting (*’-s’*) as UHGG criteria. For species with more than 1,000 genomes, we set the parameter *-g* 50000.

### Genetic distance calculation and dimensionality reduction

For each species, we computed genetic distances based on the *gene_presence_absence.Rtab* file generated by Roary. This binary matrix represents genes as rows and samples (MAGs) as columns, with values of 1/0 indicating gene presence/absence. We calculated pairwise Hamming distances between MAGs using the ‘*binary*’ method in R’s *dist()* function, generating a sample-to-sample distance matrix. Dimensionality reduction and visualization of genetic structure were performed using the *Rtsne()* function for t-SNE analysis.

### The GMR unique species-level marker genes

For species with fewer than 3,000 genomes, pan-genomes were constructed using all available genomes with Roary. For the 29 species with more than 3,000 genomes, the top 3,000 genomes, ranked by score (completeness – 5 × contamination + 0.5 × log(N50)), were selected for pan-genome construction. Core genes were identified from Roary’s *gene_presence_absence.Rtab* files, with coreness thresholds based on the number of genomes: 80% for species with fewer than 10 genomes, 60% for species with 10 to 100 genomes, and 50% for species with over 100 genomes. Species with fewer than 200 core genes were excluded from further analysis.

To identify marker genes, we employed the criteria established by MetaPhlAn4[24] for marker gene evaluation. Core genes for each species were aligned against the genomes of other species using Bowtie with the ‘*--sensitive*’ parameter. For computational efficiency, we selected up to 100 of the highest-quality genomes per species for alignment. Core genes were split into 150-nt fragments to simulate metagenomic reads, which were then mapped against representative genomes. A core gene fragment was considered a hit if it mapped to its species’ genomes above the coreness threshold and mapped to fewer than 1% of genomes from other species (quasi-markers) or not at all (perfectly unique markers). A maximum of 200 marker genes per species were selected, prioritized by uniqueness and length (longer genes first). Species with fewer than 10 markers were excluded.

This process produced a marker gene database comprising 2,777 species. We compared these species against the MetaPhlAn4 *mpa_vJun23_CHOCOPhlAnSGB_202403* database using the *mpa_vJun23_CHOCOPhlAnSGB_202403_SGB2GTDB.tsv* file. This comparison identified 579 species missing from the MetaPhlAn4 database. To create a more comprehensive resource, we integrated our marker genes with those from MetaPhlAn4 for subsequent analyses.

### Profiling microbiome composition

To assess the performance of our customized database in profiling gut microbial composition, we analyzed 1,076 publicly available fecal metagenomic samples from an independent Chinese cohort[48], 149 metagenomic samples from a German cohort[49] and 179 metagenomic samples from a Konzo cohort[50]. Taxonomic profiling was performed using MetaPhlAn4[24] (v.4.1.0) with default parameters and our custom database. The same analysis was conducted for the CGMR cohort.

## Quantification and statistical

### Comparison of microbial genomes from Chinese and non-Chinese populations

The GMR dataset contains 478,588 microbial genomes, including 247,134 from the Chinese population and 231,454 from the non-Chinese population. The median genome sizes for the two groups were 2.27Mb and 2.21Mb, respectively, with N50 lengths of 30Kb and 27Kb. Comparison between the two groups were assessed using the Wilcoxon test.

### Comparison of reads mapping

After integrating GMR-specific marker genes into the MetaPhlAn4 database, an independent Chinese cohort[48] was used for taxonomic profiling. Paired Wilcoxon tests were employed to compare the reads mapping rates between groups.

### Microbial abundance associations to human phenotypic traits

We first performed a centered log-ratio transformation on the taxonomic data and performed Spearman correlation analysis with the phenotypic data. P values were adjusted using the Benjamini–Hochberg method, and associations with an FDR-adjusted P value < 0.05 were considered significant. For species associated with multiple phenotypes, we used MaAsLin2 for multivariable association analysis with default settings, treating the phenotypes as fixed effects. MaAsLin2 was also employed to identify significant associations while adjusting for potential confounders.

### Replication of microbiome–phenotype associations

To replicate newly detected microbiome composition–phenotype associations in CGMR, we used data from 551 patients from the Qi *et al.*[25] as a replication cohort. We identified 318 associations across 20 overlapping phenotypes. To assess the variability of these associations, Cochran’s Q tests were performed to evaluate heterogeneity in effect sizes and directions between the two cohorts, using the *metagen()* function from the meta package (v.7.0.0) in R. Associations with an FDR-adjusted Cochran’s Q test P value < 0.05 were considered heterogeneous.

## Results

### Extend global gut microbial genomes with a quarter million from Chinese populations

Nearly 300,000 human gut microbial genomes were included in the late 2023 release of the Unified Human Gastrointestinal Genome (UHGG)[3] version 2, but only 30% originated from non-western populations. To enhance the representation of eastern populations and our understanding of global gut microbial genomics diversity and its health impacts, we assembled extra 6,657 gut metagenomic samples from the Chinese population [20, 25–27], which have not yet been included in the latest UHGG collection. Using a single-sample metagenomic assembly strategy (**Figure 1A**), a total of 180,719 metagenome-assembled genomes (MAGs) were obtained. We then incorporated existing microbial genomes collections, including UHGG[3], the Human Gut Microbial Biobank (hGMB)[22], the Culturable Genome Reference (CGR2)[23] and IMGG[21], with these newly assembled genomes (**Figure 1A**).

**Figure 1.**
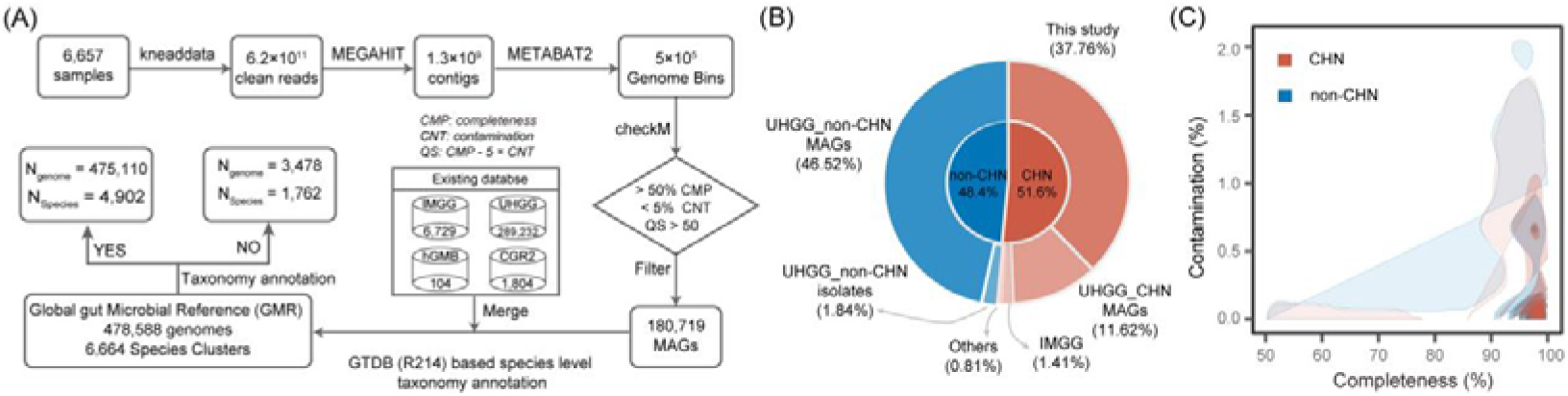
Establishment of an integrated catalog of 478,588 microbial reference genomes in the human gut. **(A)** Workflow and bioinformatic pipeline for constructing the GMR database. We assembled 180,719 high quality MAGs from 6,657 Chinese metagenomic samples with a total amount of 76.8 TB clean sequences. By integrating these MAGs with existing databases, we constructed the GMR, comprising 478,588 microbial genomes from 6,664 species, 26.4% of which are newly identified. **(B)** Summary of the microbial genomes in the GMR. The inner pie chart shows the proportion of genomes derived from Chinese (247,134 genomes, 51.6%) and non-Chinese populations. The outer pie chart highlights the distribution of genomes from different data sources. **(C)** Distribution of microbial genomes quality scores in the GMR. The X-axis represents the completeness of genomes, while the Y-axis represents contamination levels. Genomes from Chinese samples are shown in red, and those from non-Chinese samples are in blue. Color intensity reflects the number of genomes.

To standardize genome quality across all MAGs, we applied UHGG criteria[3, 51] (completeness > 50%, contamination < 5%, and quality score > 50), resulting in a total of 478,588 medium-to-high quality microbial genomes, with 247,134 (51.6%) originated from Chinese populations (**Figure 1B; Table S1**). It is worth noting that the assembled genomes were of high quality, with the median completeness and contamination rates of genomes of Chinese origin are 90.27% (interquartile range, IQR: 77.01–95.97%) and 0.88% (IQR: 0.08–1.90%), respectively. For non-Chinese genomes, these values are 90.60% (IQR: 77.47–96.51%) and 0.85% (IQR: 0.11– 1.79%), respectively (**Figure 1C; Figure S1A and S1B; Table S1**). Additionally, both median genome size (2.27Mb vs. 2.21Mb, P_Wilcoxon_=3.10×10^-35^) and N50 length (30Kb vs. 27Kb, P_Wilcoxon_=1.19×10^-109^) of Chinese-origin genomes were slightly higher than non-Chinese genomes (**Figure S1C and S1D; Table S1**). We refer to this largest integrated collection as the human Gut Microbial Reference (GMR) set. The GMR set will facilitate the decoding of global gut microbial compositional and functional diversity and its implications for regional health disparities related to the gut microbiome.

### Taxonomic classification of GMR revealed 6,664 species clusters with 1,762 being identified as potential new species

We then assessed the taxonomic distribution of microbial genomes in the GMR using a two-step approach. In brief, we first annotated all 478,588 genomes at the species level using the Genome Taxonomy Database (GTDB, R214)[35, 36]. Among these, 475,110 genomes were successfully assigned to 4,902 species. For the remaining 3,478 genomes that could not be annotated at the species level, we clustered them using dRep[37] with an ANI threshold of > 95%, resulting in 1,762 clusters (**Figure 2A; Table S2**). Of these 1762 clusters, 478 contained at least 2 genomes (**Figure S2**). Notably, among the 1,762 clusters that could not be assigned to known species, 101 clusters (1.52%) and 4 clusters (0.06%) could not even be classified at the genus or family levels, respectively. In total, we obtained 6,664 species-level clusters, comprising 6,624 bacterial and 40 archaeal species (**Figure 2A; Table S2**). The most prevalent bacterial phyla were *Bacillota_A* (2,585 species), *Actinomycetota* (1,103 species), and *Bacteroidota* (929 species) (**Figure 2A**). Among the phylum with more than 2 species, *Patescibacteria* has the highest proportion (43.48%) of unknown species discovered (**Figure 2B**). This highlights the vast, unexplored microbial diversity within the human gut.

**Figure 2.**
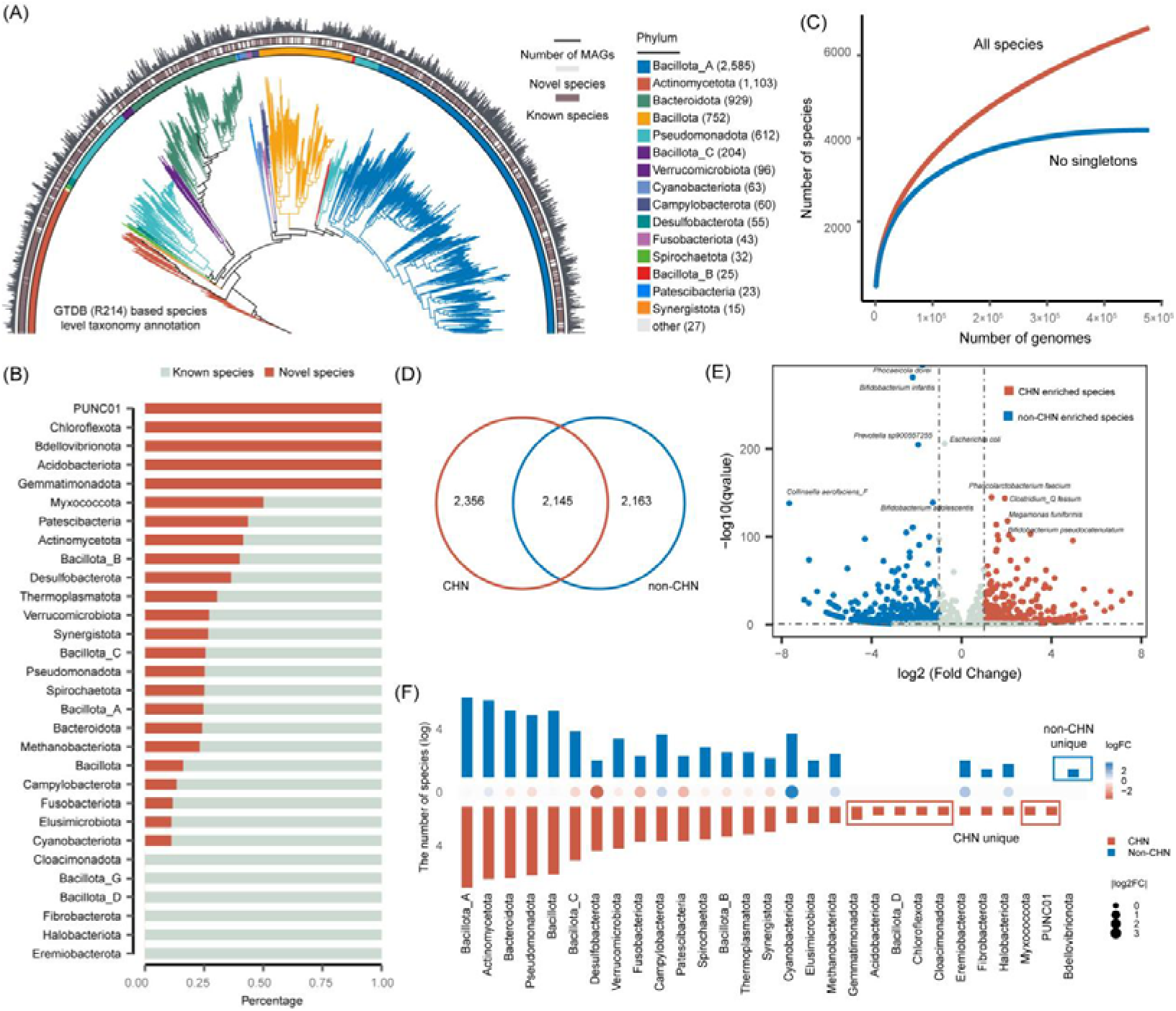
Taxonomic classification of the GMR reveals 6,664 species clusters with 1,762 identified as potential novel species. **(A)** Phylogenetic tree of all 6,624 bacterial species detected in the human gut. The colors on the branches represent the bacterial phyla, while the color on the outer circle indicates newly identified species based on GTDB R214 database. The length of the outer bars represents the total number of MAGs obtained for each representative species. **(B)** The proportion of newly characterized species within each phylum, with red indicating the newly identified species. **(C)** Rarefaction curves of the number of species detected as a function of the number of nonredundant genomes analyzed. Curves are depicted both for all the GMR species and after excluding singleton species (represented by only one genome). **(D)** Overlap of microbial species clusters between Chinese and non-Chinese populations. In general, 2,145 species have been found to be presented in both Chinese and non-Chinese populations. **(E)** Comparison of 2,145 shared species between Chinese and non-Chinese populations. Among them, 896 species (41.77%) exhibited significant population enrichment. The X-axis represents the ratio of the number of genomes in the Chinese population to that in the non-Chinese population, expressed as log2. The Y-axis represents the P-value from the chi-square test with multiple corrections. Each dot represents a species, with red indicating species enriched in the Chinese population and blue indicating species enriched in the non-Chinese population. **(F)** Distribution of population-specific species in Chinese and non-Chinese populations. Red represents species unique to the Chinese population, and blue represents species unique to the non-Chinese population. The X-axis indicates phylum classification, and the Y-axis represents the number of species, displayed on a log scale.

The rarefaction curve analysis suggests that the currently detected microbial species have not yet reached a plateau, indicating that a substantial number of undiscovered species may still exist in the human gut microbiome (**Figure 2C**). However, these potential novel species are likely to be rare members of the gut microbial community, as supported by considering only species with at least two conspecific genomes, the species accumulation curve approaches saturation (**Figure 2C**).

Species distribution analysis based on the GMR database identified a total of 4,501 gut microbial species in the Chinese population and 4,308 species in non-Chinese populations, with 2,145 core species shared between the two groups (**Figure 2D; Table S2**). Notably, among these globally prevalent core species, 896 species exhibited significant population-specific enrichment (Chi-squared test, FDR < 0.05) (**Figure 2E; Table S3**). For instance, *Collinsella aerofaciens_F*, *Alistipes excrementigallinarum* and *Phocaeicola sp000432735* showed a marked genomic abundance in non-Chinese populations, whereas *Collinsella sp003439125*, *Haemophilus_D sp905205995*, and *Collinsella sp003459245* were significantly enriched in the Chinese population (**Figure 2E; Table S3**).

Further analysis revealed that 2,356 gut microbial species were unique to the Chinese population, predominantly belonging to the phyla *Gemmatimonadetes*, *Acidobacteria*, *Firmicutes_D*, *Chloroflexi*, *Calditrichaeota*, *Myxococcota*, and *PUNC01* (**Figure 2F**). Notably, *Bdellovibrionota* was undetectable in all Chinese genomes, which may be attributed to specific environmental factors and dietary habits in China that are unfavorable for *Bdellovibrionota* growth, or limitations in sampling scope and depth. This finding provides crucial insights into population-specific differences in the human gut microbiome and their underlying mechanisms.

### Unveiling functional potential and resistome dynamics in the human gut microbiome

To systematically characterize the functional potential encoded with microbial genomes in our GMR set, we performed comprehensive protein-coding sequence (CDS) prediction across all 478,588 assembled genomes. This analysis identified 1,028,234,783 putative genes, averaging 2,148 CDS per genome (**Figure 3A; Table S4**). To comprehensively elucidate the functional diversity of the gut microbiota, we annotated the proteins using currently available databases, including UniProt, RefSeq, Pfam, TIGRFAM and ResFinder. Overall, 47% of the genes could be matched to at least one database. The median proportion of genes with unknown functions was 47.3% (IQR: 43.3–51.5%) in Chinese genomes and 48.2% (IQR: 43.7–52.9%) in non-Chinese genomes (**Figure 3B**). This indicates that nearly half of the microbial genes in the human gut remain unexplored (**Figure 3C**), reinforcing previous estimates[4, 52] of microbial “dark matter”. Notably, we found that, on average, species in the *Thermoplasmatota* phylum had the highest percentage of unknown functional genes (**Figure S3**), suggesting this lineage represents a particularly promising target for functional discovery.

**Figure 3.**
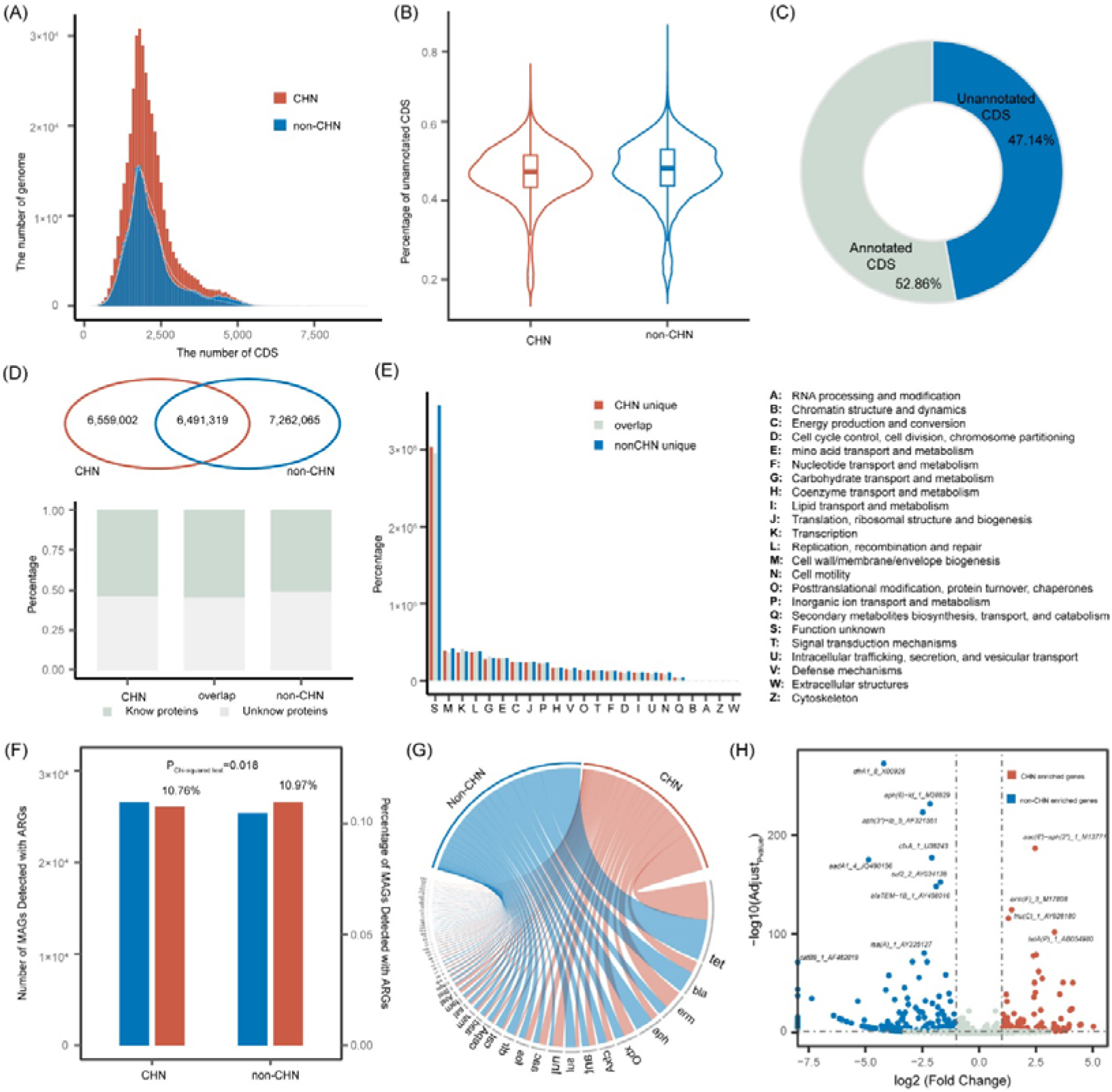
Functional capacity of the human gut microbiota. **(A)** Distribution of the total number of CDSs (coding sequences) per microbial genome from Chinese and non-Chinese populations. On average, genomes from Chinese and non-Chinese populations contained 2,126 and 2,172 CDSs per genome, respectively. **(B)** Comparison of un-annotatable CDS percentages between Chinese and non-Chinese populations. Boxplots display the median values (center lines), interquartile ranges (IQRs, represented by the box edges), and whiskers, which extend to the most extreme data points within 1.5 times the IQR from the lower and upper quartiles. **(C)** Pie chart showing the proportion of functionally annotated versus unannotated CDSs. Based on the UniProt, RefSeq, Pfam and TIGRFAMs databases, 47.14% of CDSs remain unannotated. **(D)** Distribution of encoded proteins clustered at 90% amino acid similarity between Chinese and non-Chinese populations. A total of 6,491,319 protein clusters are shared by both groups. The bars at the bottom indicate the number of proteins encoded by cluster representatives from three categories (unique to Chinese, overlap, and unique to non-Chinese), stratified into known and unknown protein clusters. **(E)** COG functional annotation of the GMRP catalog clustered at 90% amino acid identity. Only functions with >5000 genes are plotted. **(F)** Distribution of antibiotic resistance genes (ARGs) detected in MAGs from Chinese and non-Chinese populations. Blue bars represent the number of MAGs with at least one ARG detected, while red bars indicate the proportion of MAGs containing ARGs relative to the total number of MAGs in each population. **(G)** Visualization of antibiotic resistance genes (ARGs) detected in the human gut microbiome. **(H)** Population enrichment of detected antibiotic resistance genes (ARGs). The X-axis represents the ratio of the number of genomes with ARGs in the Chinese population to that in the non-Chinese population, expressed as log2. The Y-axis represents the P-value from the chi-square test with multiple corrections. Each dot represents a gene, with red indicating genes enriched in the Chinese population and blue indicating genes enriched in the non-Chinese population.

Given the high complexity and vast number of genes identified in the GMR, clustering analysis was employed to group functionally similar genes, thereby facilitating the identification of key functional categories and their potential roles in the ecosystem. Subsequent clustering of these gene products at 50%, 90%, and 100% amino acid identity thresholds yielded 7-200 million protein clusters (**Figure S4**) (We termed GMRP50, GMRP90 and GMRP100, respectively). Comparative analysis of GMRP90 clusters revealed distinct features between Chinese and non-Chinese populations. While 6,491,319 protein clusters were shared between populations, population-specific clusters were identified (6,559,002 unique to Chinese, 7,262,065 to non-Chinese) (**Figure 3D**), indicating potential biogeographic specialization of gut microbial functions. On the basis of the distribution of COG (Clusters of Orthologous Genes) functions, core microbial processes were revealed, with cell wall/membrane biogenesis (5.81%), transcription (5.57%), and replication/recombination/repair (5.66%) being the most functional categories (**Figure 3E; Table S5**).

Furthermore, our analysis extended beyond basic physiology to address critical public health concerns. Given the significant public health threat posed by antibiotic resistance genes (ARG) and the role of microorganisms as efficient vectors for their dissemination, we examined the distribution differences of ARGs between Chinese and non-Chinese populations. Among 51,995 MAGs, a total of 1,050 ARGs were detected across 2,233 species (**Figure S5; Table S6**), indicating that approximately 33.5% of gut microbes harbor resistance genes—underscoring the severity of the antibiotic resistance challenge. Strikingly, one MAG from a Chinese individual, classified as *UBA7597 sp900542935*, harbored 38 antibiotic resistance genes, the highest observed number (**Table S7**). At the species level, *Escherichia coli*, *Klebsiella pneumoniae*, *Phocaeicola dorei*, *Enterococcus_B faecium* and *Bacteroides uniformis* were the top five species with the highest number of resistance genes (**Figure S5; Table S6**).

Furthermore, among the 51,995 ARG reservoirs MAGs, 26,594 were derived from Chinese individuals and 25,401 from non-Chinese individuals, with the resistance gene detection rate slightly higher in the non-Chinese population (10.97%) compared to the Chinese population (10.76%) (Chi-squared test, P=0.018) (**Figure 3F**). *Tet* is the most frequently detected antibiotic resistance gene (**Figure 3G; Table S8**). Of the 1,050 resistance genes, 195 were exclusively detected in Chinese genomes and 311 exclusively in non-Chinese genomes, with some genes showing marked population-specific enrichment (**Figure 3H; Table S8**). For example, *dfrA1*, *aph(6)* and *aph(3)* were significantly enriched in non-Chinese population, while *aac(6’)-aph(2”)*, *erm(F)* and *tetA(P)* were significantly enriched in Chinese population (Chi-squared test, FDR < 0.05) (**Figure 3H; Table S8**). Moreover, although some species were common to both populations, certain resistance genes were detected only in one group. For instance, the *dfrA8* and *dfrA7* genes were exclusively found in *Escherichia coli* MAGs from non-Chinese genomes, *tmexD2* was detected only in *Pseudomonas aeruginosa* MAGs from non-Chinese genomes, and *cfiA3* was exclusively observed in *Bacteroides fragilis_A* MAGs from Chinese genomes (**Table S8**).

Consequently, the GMR set not only significantly expands the current microbiome database but also provides a comprehensive catalog of gut microbiome proteins, offering a valuable resource for decoding global gut microbial functional diversity and exploring its implications for regional health disparities and future functional metagenomic analyses.

### Genetic stratification of predominant species between populations is related to antibiotic resistance

For 2,145 out of 6,664 species that present in both Chinese and non-Chinese populations (**Figure 4A**), we questioned their potential genomic differences and underlying functionalities. For this, we focused on 304 species with at least 100 MAGs and a proportion of at least 30% in either Chinese and non-Chinese populations (**Figure 4A and 4B**). Notably, 58.88% of these species belonged to the phylum *Bacillota_A* (**Figure S6**), with genome counts ranging from 100 to 11,423, totaling 306,525 MAGs (**Figure 4B; Table S9**). For each species, we conducted a pan-genome analysis, assessing the presence or absence of each gene within the species. Based on the distribution of pan-genes, we calculated intra-species genetic distances. Subsequently, we employed t-distributed Stochastic Neighbor Embedding (t-SNE) to characterize genetic structures and explore their population-specific differences. Strikingly, we found that 225 species (74%) exhibited significant genetic structural divergence between Chinese and non-Chinese populations (**Figure 4C; Table S10**). Among these, species from the *Lachnospiraceae* (24.4%), *Bacteroidaceae* (13.78%), and *Ruminococcaceae* (8%) families were significantly enriched (Hypergeometric test, P = 3.82 × 10^-12^, P = 7.62 × 10^-7^ and P = 7.61 × 10^-4^ respectively) (**Figure 4D, Table S11**), suggesting a potential population-specific adaptation in these taxa.

**Figure 4.**
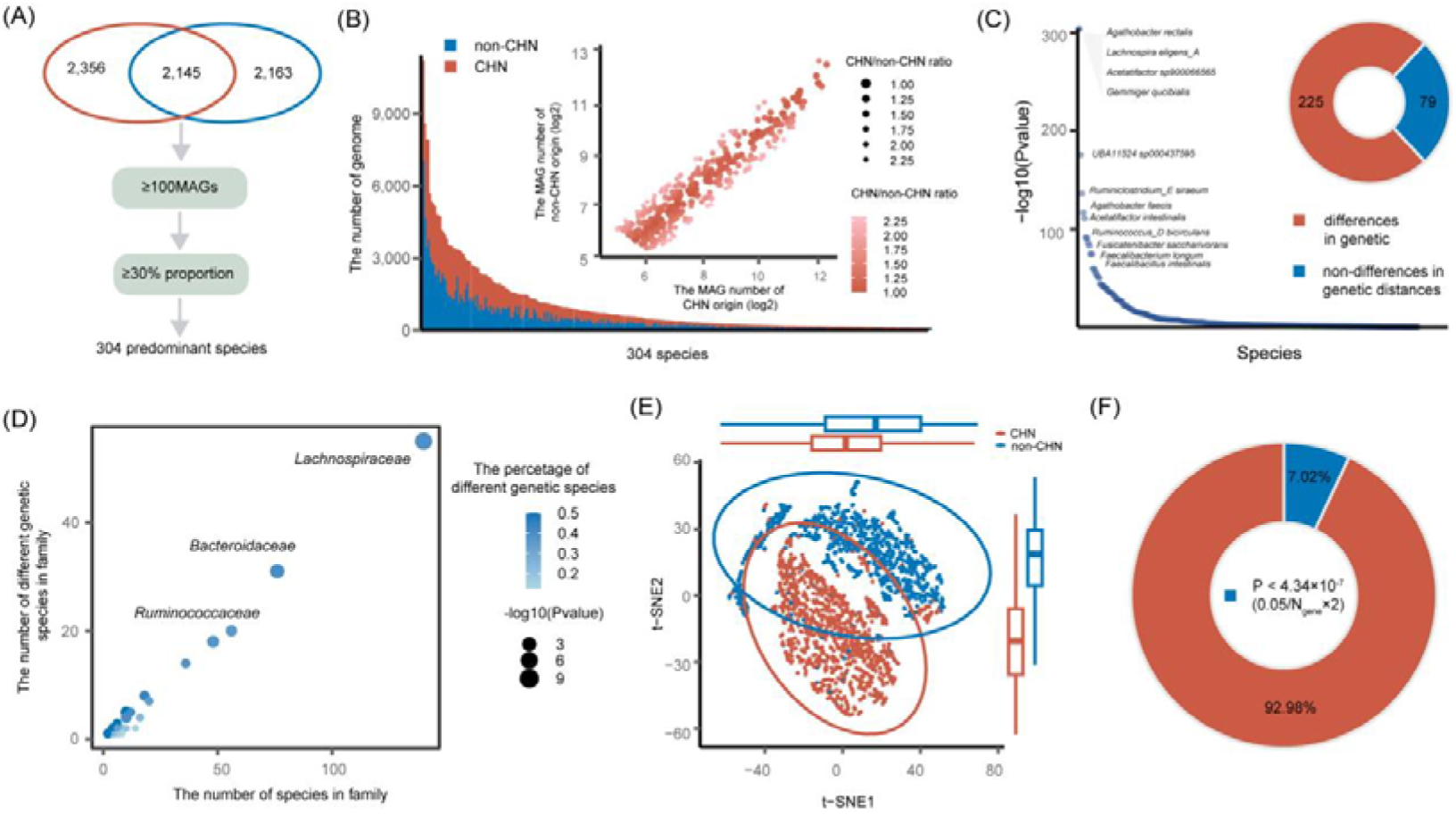
Genetic stratification of predominant species between populations. **(A)** Distribution of predominant species in the GMR. A total of 2,145 species were shared between Chinese and non-Chinese populations, among which 304 species showed comparable abundances (with >100 MAGs in both groups). **(B)** Summary of MAGs per species clusters from Chinese and non-Chinese populations. The bar plot on the left depicts the 304 species clusters with at least 100 MAGs and a proportion of at least 30% MAGs in either Chinese and non-Chinese populations. The scatter plot on the right shows the proportion of MAGs per species cluster from Chinese and non-Chinese populations. The darkness of color gradient and the size of the dots indicate proportions. **(C)** Population-specific genetic variation in 225 microbial species (represented in the right circos plot). The dot plot displays –log10 transformed p-values for species association with tSNE coordinates, with the top 12 most significant species labeled. **(D)** Summary of microbial species with significant genetic variation between populations at family level. The X-axis represents the total number of species within each family, while the Y-axis shows the number of species showed significant genetic stratification between Chinese and non-Chinese populations. Each dot represents one family with the size of the dots indicates P-value from Hypergeometric test, and the darkness of color represents the proportion of species showed significant stratification between Chinese and non-Chinese populations within each family. Among the 45 families examined, only three families exhibited a significant correlation at an FDR threshold of less than 0.05 and was subsequently labeled with its corresponding family name. **(E)** Population-level genetic structure of *Lachnospira eligens_A*. The scatter plot represents tSNE coordinates, with corresponding boxplots showing distribution patterns along each tSNE axis. **(F)** Identification of 4,044 genes showing significant associations with genetic structure. The significance threshold was set at 0.05/(total gene number × 2 tSNE dimensions) after multiple testing correction.

Given its importance in previous studies, *Prevotella copri* serves as a reference for understanding the population structuring of gut species. Previous studies have confirmed that *P. copri* exhibits a genetic structure linked to geographic origin and lifestyle, with Chinese-origin genomes displaying distinct lineages [15–18]. Consistent with this, we observed obvious genomic differences between Chinese and others (**Figure S7; Table S10**). Besides this, the most distinct population-level genetic stratifications were observed in species such as *Agathobacter rectalis*, *Lachnospira eligens_A*, *Acetatifactor sp900066565*, and *Faecalibacillus intestinalis* (**Figure 4C; Table S10**). For example, *Lachnospira eligens_A* comprised 4,818 genomes in the GMR database, with 2,519 originating from Chinese individuals and 2,299 from non-Chinese individuals. The genetic structure of *L. eligens_A* differed significantly between the two populations (**Figure 4E**), with phylogenetic analysis revealing that Chinese-derived genomes formed a distinct clade (**Figure S8**). To further investigate the drivers of this genetic differentiation, we performed an association analysis between the presence/absence of 57,644 pan-genome genes and the t-SNE genetic clustering. We identified 4,044 genes (7.02%) that significantly contributed to the observed genetic stratification (**Figure 4F**). By further annotating these genes, we observed a wide range of encoded functionalities, from glycolipid metabolism to antibiotic resistance. For instance, we observed that the presence or absence of antibiotic resistance–associated genes, such as the methicillin resistance regulator mecI and the multidrug efflux pump *mepA*, significantly contributed to the population-specific genetic structure of *L. eligens_A* (**Figure S9**). This suggests that variations in antibiotic resistance among strains from different regions may reflect distinct selective pressures in the respective populations. Therefore, potential differences in resistance profiles should be carefully considered when designing functional experiments involving this species.

### Taxonomic markers from new species boost populational profiling by up to 23%

For the 1,762 newly assembled species, to enhance the detection of these microbes in population studies and provide a more comprehensive gut microbial composition profile, we integrated MAGs from our GMR set into the MetaPhlAn4 taxonomic classification database[24], released in March 2024. Among the 6,664 species-level clusters, 2,465 are represented by a single genome, and 929 by two genomes (**Figure 5A**; **Figure S10; Table S9**). For the remaining 3,270 species clusters with at least three genomes per cluster, we classified their species-level specific markers.

**Figure 5.**
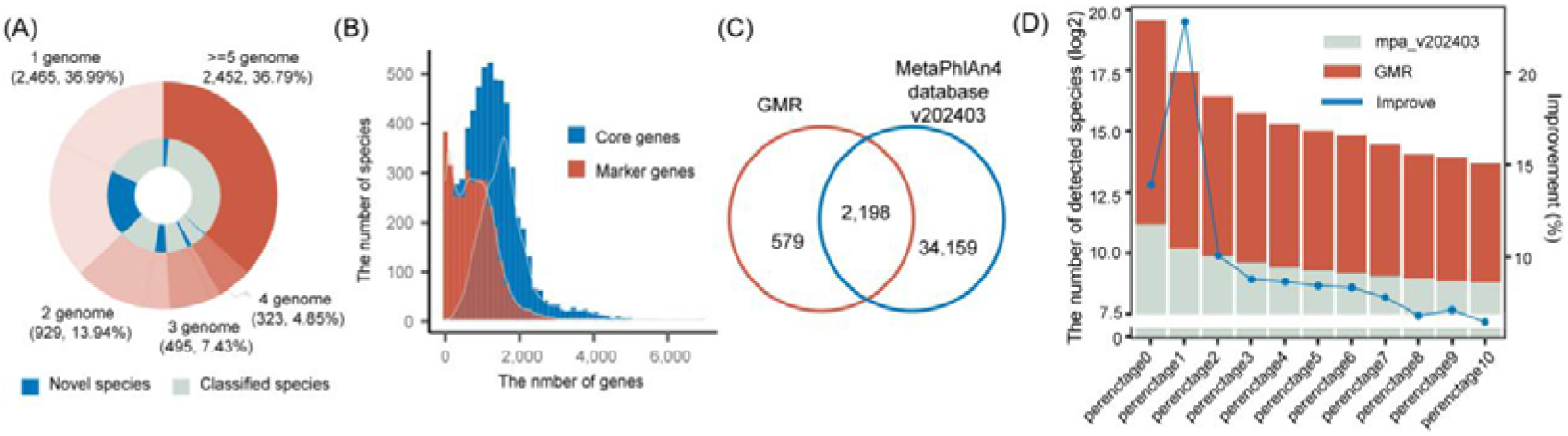
Integrating microbial genomes from newly classified species into MetaPhlAn4 enhances taxonomic profiling. **(A)** Distribution of microbial genomes across species-level clusters. The inner circle represents the proportion of novel and classified species within each level. A total of 3,270 species clusters contains at least 3 genomes. **(B)** Summary of core and marker genes identified in each species. On average, each species contains 1,626 core genes and 676 marker genes. **(C)** Overlap between GMR species (with at least 10 species-specific marker genes) and species in the MetaPhlAn4 database. The GMR contributes additional species-specific marker genes for 579 species, complementing the MetaPhlAn4 database. **(D)** Taxonomic profiling at the species level using the newly constructed database in an independent Chinese cohort with diverse filtering cutoffs. A total of 1,076 publicly available fecal metagenomic samples were analyzed. The X-axis shows different filtering cutoffs from 0% to 10%. The left Y-axis (bar chart) represents the number of detected species, with the red portion highlighting the contribution of newly identified species in the GMR. The right Y-axis (blue line) represents the improvement in species detection achieved by including the newly identified species from the GMR.

Following the MetaPhlAn4 taxonomic database construction pipeline, we first identified core coding sequences (CDS) for each species, averaging 1,626 core genes per species (**Figure 5B**; **Table S12**). We then screened these core genes to identify species-specific marker genes, finding an average of 676 per species (**Figure 5B**; **Table S12**). Based on the MetaPhlAn4 guidelines, we retained a maximum of 200 marker genes for each species. For 493 species with fewer than 10 species-specific marker genes and 2,198 species overlapping with the existing MetaPhlAn4 taxonomic database, we excluded them from the database construction (**Figure 5C)**. Ultimately, 96,156 marker genes from 579 non-overlapped species were integrated into the MetaPhlAn4 database to build our customized database (**Table S13**).

To evaluate the performance of our customized database in profiling gut microbial composition, we analyzed an independent cohort of 1,076 publicly available fecal metagenomic samples from a Chinese cohort[48]. Using MetaPhlAn4 with default parameters and our customized taxonomic classification database, we performed microbial abundance profiling. On average, integrating the GMR database increased the reads mapping rate by 1% per sample (**Figure S11**). In total, we detected 2,669 species across all samples, including 326 newly added species. Filtering for species present in at least 1% of samples, 150 out of 1,325 species were from our new additions, indicating a 22.8% increase compared to the existing MetaPhlAn4 database (**Figure 5D**). With a filtering threshold set from 1% to 10%, our customized database consistently showed an average 9.5% improvement in the total number of species detected. The species that showed an increased detection rate at the family level primarily belong to *Acutalibacteraceae*, *CAG-508* and *Ruminococcaceae*, as well as 13 other families in total (Hypergeometric test, FDR < 0.05) (**Figure S12; Table S14**). Notably, 97.58% of individuals had at least one newly added species detected, indicating that these species are widely present in Chinese populations (**Figure S13**).

To further validate the application value of the constructed personalized database in non-Chinese populations, we conducted species abundance characterization analyses using two completely independent datasets: one comprising 149 German (European) samples[49] and the other comprising 179 Congolese (African) samples[50], none of which were involved in the database construction. In the European cohort, 1,895 species were identified, with 222 (13.27%) contributed by the GMR database compared to MetaPhlAn4. When the abundance threshold increased from 1% to 10%, the GMR database provided an additional 8.79% contribution on average (**Figure S14**). In the African cohort, 1,563 species were detected, with 206 (15.18%) contributed by GMR. After filtering species present in <1% of samples, 170 of the remaining 1,304 species originated from GMR; at a 10% threshold, 60 of 601 species were GMR-derived, with an average contribution of 13.41% (**Figure S15**). These results highlight the enhanced capability of our customized taxonomic classification database for species abundance identification in global microbiome studies.

### Abundance of newly profiled species widely associated with host phenotypic traits

Next, we assessed the potential biological significance of these newly profiled species in the CGMR cohort[20]. Using our customized database, we detected 5,310 species in 3,234 participants, including 465 newly added species (**Figure S16; Table S15**). On average, 201 species and 11 new species were detected per sample (**Figure 6A**). Unsurprisingly, 97.8% of individuals had at least one newly added species detected (**Figure 6A; Table S16**).

**Figure 6.**
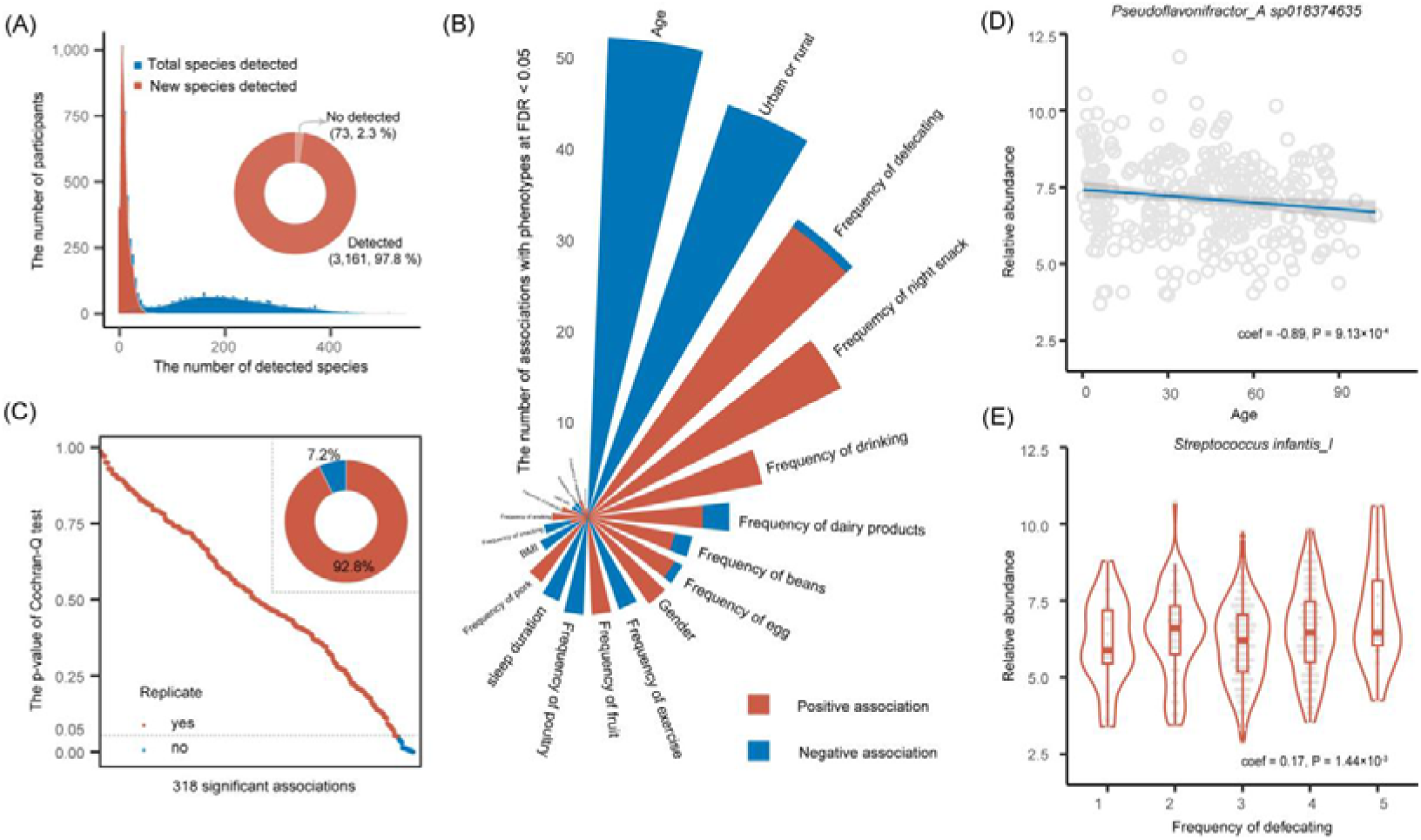
Abundance of newly classified species is widely associated with host intrinsic traits and dietary habits across independent cohorts. **(A)** Summary of microbial species profiled in a cohort of 3,234 participants. The blue bars represent the total number of detected species, while the red bars indicate the number of newly classified species in the GMR. On average, 201 species were detected per sample, with 11 belonging to newly classified species in the GMR. The pie chart on the right illustrates the proportion of samples with at least one newly detected species, showing that 97.8% of the samples contained at least one new species. **(B)** Host phenotypic associations with the abundance of newly profiled species. Bars represent different phenotypic traits, sorted in descending order by the number of significant associations. The direction of the associations is color-coded: red indicates positive associations, while blue represents negative associations. The length of each bar reflects the number of significant associations (FDR < 0.05). **(C)** Replication of 318 significant associations in an independent cohort of 551 participants. The Y-axis shows P-values from the Cochran-Q test for phenotypic associations with microbial species, while the X-axis displays the 318 associations ordered by heterogeneity test P-values. The pie chart in the upper right shows the proportion of replicable associations, with blue indicating P-values < 0.05 and red indicating P-values > 0.05 in the Cochran-Q test. **(D)** The relative abundance of *Pseudoflavonifractor_A sp018374635* was negatively associated with age. The X-axis shows participant ages, and the Y-axis represents species abundance. The solid line represents linear regression, with gray shading indicating the 95% confidence interval. Each dot represents a participant. **(E)** The relative abundance of *Streptococcus infantis_I* was significantly higher with increased defecation frequency. The box plot compares species abundance across different defecation frequencies. The boxes represent the median, first, and third quartiles (25th and 75th percentiles), with whiskers extending to the most extreme data points within 1.5 times the interquartile range. Each dot represents a participant.

By using spearman correlation to associate the abundance of 54 out of the 465 newly added species with 22 phenotypic traits (each with at least 5% prevalence), we identified 325 significant associations (FDR < 0.05, **Figure 6B; Table S17**), ranging from intrinsic factors to diverse dietary habits. Among them, age showed the most associations with the abundance of these novel species (**Figure S17**). To assess the robustness of these associations, we used an independent cohort consisting of 551 participants[25]. We identified 318 associations for 20 overlapping phenotypes (**Table S18**). Heterogeneity analysis further revealed that 295 out of 318 (92.8%) associations were consistent between the two cohorts (P_Cochran-Q_ > 0.05) (**Figure 6C; Table S18**).

We then assessed the potential confounding effects of the identified associations, noting that multiple phenotypic traits were associated with the same species. Using MaAsLin2[53], a tool for addressing multivariable associations between host phenotypes and gut microbes, we identified 64 robust associations between 39 species and 16 phenotypes among those 295 associations (**Table S17**). These associations are primarily related to urban or rural residency, defecation frequency, age, lifestyle, and dietary habits. For example, the abundance of *Pseudoflavonifractor_A sp018374635* was negatively associated with age (P = 9.13×10^-4^, **Figure 6D**), while there was a positive association between defecation frequency and the abundance of *Streptococcus infantis_I* (P = 1.44×10^-3^, **Figure 6E**). Additionally, we observed a significantly higher abundance of a *Streptococcus s*pecies *GT_GMRSGB00117* in urban areas compared to rural areas (P = 1.44×10^-2^, **Figure S18A; Table S17**). Regarding lifestyle, the frequency of drinking was positively associated with the abundance of *Nanosynbacter sp900556355* (P = 2.47×10^-2^) (**Figure S18B; Table S17**).

These results demonstrate that our customized database can uncover previously undetected relationships between microbes and host phenotypic traits, laying the foundation for further exploration of the relationship between microbial abundance and host disease.

## Discussion

Studies have shown that the composition and functionality of gut microbiota vary markedly between individuals[2, 54, 55]. Given that the Chinese population accounts for over 15% of the global population, key questions remain: What are the defining features of the gut microbiota in Chinese individuals, and how do these features differ from those in Western populations? Although several studies[20, 21] have analyzed the gut microbiota of Chinese cohorts, many are limited either by focus on specific ethnic groups or by small sample sizes, thus failing to fully capture the unique characteristics of the Chinese gut microbiome. In this study, we established the most comprehensive human gut microbial resource database to date—the Global Microbiome Reference (GMR)—which comprises 478,588 high-quality microbial genomes, including 247,134 from Chinese individuals and 231,454 from non-Chinese populations. This database contains over 50% more genomes than the UHGG and effectively addresses the underrepresentation of non-Western microbial genomes, thereby balancing the dataset between Chinese and Western populations. Notably, these genomes demonstrate high consistency across critical quality metrics (genome length, N50, completeness, and contamination), providing a robust foundation for comparative analyses of species composition and functional profiles. Based on the GTDB R214 database, these genomes are classified into 6,664 species-level clusters, with 1,762 identified as potential novel species. Globally, 2,145 species are shared between Chinese and non-Chinese populations, whereas 2,356 and 2,163 species are unique to the Chinese and non-Chinese populations, respectively—differences that underscore the distinct microbial distribution among diverse human groups.

It is particularly noteworthy that certain phyla—such as *Gemmatimonadota*, *Bacillota_D*, *Acidobacteriota*, *Chloroflexota*, *Myxococcota*, *Cloacimonadota*, and P*UNC01*—appear exclusively in Chinese individuals, while *Bdellovibrionota* has not been detected in the Chinese gut. Additionally, for species shared between populations, there is marked population-specific enrichment. For example, *Collinsella sp003439125* is represented by 181 metagenome-assembled genomes (MAGs) in the Chinese cohort but only 1 in non-Chinese populations, whereas *Collinsella aerofaciens_F* is present as 610 MAGs in non-Chinese individuals compared to just 3 in Chinese individuals. Although geographic, climatic, dietary, and lifestyle factors undoubtedly influence the distribution and abundance of specific microbial taxa[55–59], further systematic studies are needed to determine the extent to which these factors, as opposed to sampling limitations, drive these differences.

Regarding the protein sequences encoded by the GMR genomes, we conducted extensive functional annotation using all available databases—including Prokka, COG, GO, and KEGG. Despite these comprehensive efforts, approximately 47% of the protein sequences remain functionally uncharacterized, representing a significant fraction of genomic “dark matter.” This gap highlights our limited understanding of the long-term residents of the human gut microbiome and calls for additional functional studies.

Our analysis also revealed that the detection rate of antibiotic resistance genes (ARGs) is higher in non-Chinese populations than in Chinese populations, with tet genes being the most prevalent. Moreover, we identified a “superbug” genome that harbors 38 ARGs, underscoring the need for caution in antibiotic treatment strategies and emphasizing the potential public health risks associated with multidrug resistance.

The results of our study indicate that species unique to the Chinese population constitute over 35% of the total microbial diversity. Excluding these species from current reference databases would likely bias microbiome analyses in Chinese cohorts and may even lead to the omission of key signals related to host physiology and disease. MetaPhlAn (Metagenomic Phylogenetic Analysis), a classic marker gene-based tool known for its efficiency and scalability, has become a cornerstone in microbiome research[24, 60, 61]. By integrating 579 species unique to the GMR into the existing MetaPhlAn database, we developed an updated version that is more representative and comprehensive. When applied to independent cohorts from China, Europe, and Africa, this updated database increased species detection rates by an additional 9.5%, 8.75%, and 13.41%, respectively. This significant improvement not only demonstrates the advantage of incorporating Chinese-specific microbial species but also provides a richer dataset for elucidating the complex relationships between the microbiome and host health.

Despite these advances, our study has several limitations. First, our comparative analysis focused solely on the differences between Chinese and non-Chinese populations in terms of species abundance and gene function, without a systematic evaluation of microbial variations across different countries and geographic regions. This limitation may constrain our broader understanding of gut microbiota diversity among various ethnic groups. Second, our investigation of species and functions unique to Chinese and non-Chinese populations remains preliminary; we have not yet explored the underlying causes of these differences or their impacts on disease phenotypes. Third, even though we have incorporated 579 new species into the MetaPhlAn database, the limitations in sample size and data sources may still result in the omission of low-abundance or rare taxa. Furthermore, the validation of our expanded database relied primarily on cohorts from Europe, Africa, and China, which have relatively limited sample sizes. Future research should include larger and more diverse datasets from additional regions to ensure the global applicability and robustness of the updated database. Finally, while our correlation analyses suggest associations between newly identified species and host phenotypes, experimental validation is necessary to confirm the specific roles these microbes play in host physiology—a critical direction for future research.

The establishment of this comprehensive and representative GMR collection is a pivotal step toward advancing our understanding of the human gut microbiome’s role in health and disease. By addressing the existing biases inherent in earlier databases, our study enables a more accurate and inclusive depiction of microbial diversity across different ethnic and geographic groups. Such advancements are essential for developing targeted diagnostic tools and therapeutic strategies that are sensitive to population-specific microbiome profiles. Ultimately, our work not only fills a critical gap in current microbial genomic resources but also paves the way for future investigations into the complex interactions between the gut microbiota and host physiology. This integrated approach promises to enhance the precision of microbiome-related research and offers new avenues for personalized healthcare interventions, ensuring that discoveries in microbial ecology are translated into meaningful clinical applications.

### Conclusions

In this study, we established a globally representative and population-balanced gut microbial genome reference – GMR – by assembling over 478,588 high-quality genomes from Chinese and non-Chinese populations. Our work reveals substantial compositional and functional differences in the gut microbiome across populations, with over one-third of species being population-specific and 74% of prevalent species showing genomic stratification. The discovery of millions of novel genes, many of unknown function, underscores the untapped microbial diversity in underrepresented populations. By enhancing existing MetaPhlAn4 with our dataset, we achieved significantly improved profiling resolution and identified robust microbiome-host trait associations. This resource lays a critical foundation for future studies into microbiome-driven mechanisms underlying health and disease, and emphasizes the importance of including diverse populations in microbiome research to address global health disparities.

### Lead contact

Further information and requests for resources and reagents should be directed to the Lead Contact lianmin Chen.

### Data and code availability

The 478,588 mircobial genomes of GMR have been uploaded to the National Genomics Data Center (NGDC) with the accession number OMIX006649 (https://ngdc.cncb.ac.cn/omix/release/OMIX006649). The metabolome data of isolates are deposited in the National Genomics Data Center (NGDC) with accession number OMIX004554. The custom MetaPhlAn4 database can be download from https://zenodo.org/records/13953612. The workflow used to generate the MAGs from metagenomic data and statistical analysis codes are available via https://github.com/MicrobiomeCardioMetaLab/GMR_project.

## Acknowledgements

This project was funded by the National Natural Science Foundation of China (NSFC, 32394052, 32270077 and Excellent Young Scientists Fund Program Overseas-2022); Jiangsu Shuangchuang Project (JSSCTD202450); the Natural Science Foundation of Jiangsu (BK20220709); High-level Talent Cultivation Program (CZ0121002010039 & YNRCZN0301) and Major Basic Research Fund (TS202403) of Jiangsu Province Hospital; Changzhou Medical Center (CMCM202204); Development of Jiangsu Higher Education Institutions Priority Academic Program (PAPD). The funders had no role in the study design, data collection and analysis, decision to publish, or preparation of the manuscript.

## Author contributions

L.C, Y.Z., X.K., and Q.Z. contributed to conceptualization and funding. The CGMR Consortium Initiative staffs contributed to data and sample collection. Q.D., B.M., X.Z. and L.C. contributed to data analysis. Q.D., L.C., Y.Z., X.K., Q.Z., B.M., X.Z., and P.H. drafted the manuscript. M.G., S.Y., Y.J., Y.Z., Z.S., Q.D., D.H., X.W., L.L., C.Z., C.Z., M.K., C.H., T.W., H.Z., J.S., Y.S., Y.W., L.T., S.H., H.Z., W.S., and W.C. contributed to discussion of the content. All authors read and approved the final manuscript.

## Figure Legends

**Figure S1.**
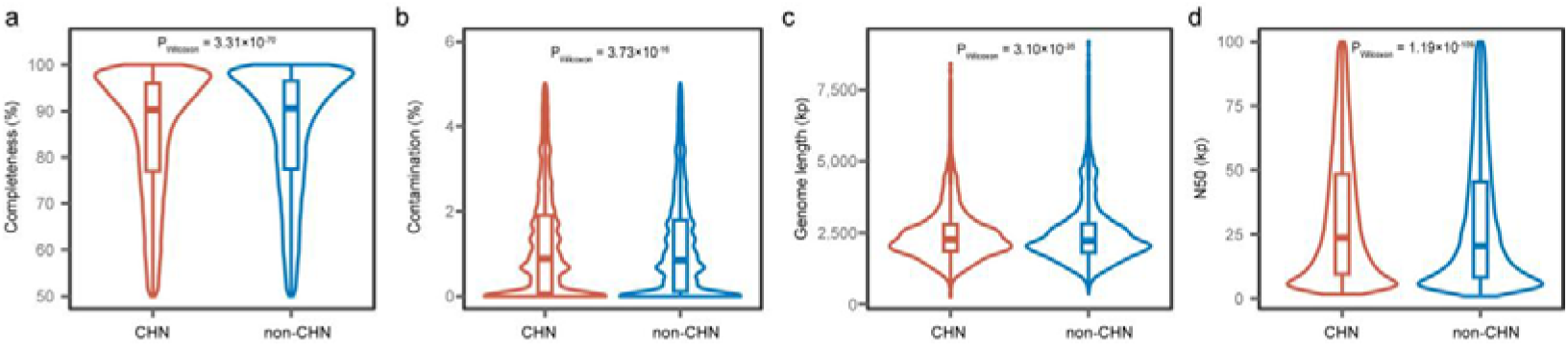
Summary of microbial genomes in the GMR from Chinese and non-Chinese populations. Panels show the distribution of completeness (**A**), contamination (**B**), genome length (**C**), and N50 length (**D**) across the 478,588 genomes included in the GMR. The boxplots display the median values (center lines), interquartile ranges (IQRs, represented by the box edges), and whiskers, which extend to the most extreme data points within 1.5 times the IQR from the lower and upper quartiles.

**Figure S2.**
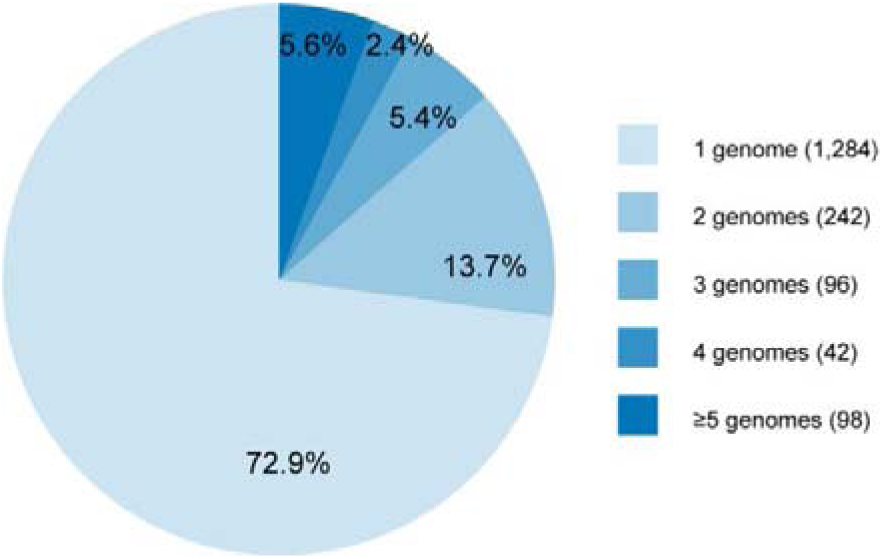
Distribution of microbial genomes across 1,762 novel species-level clusters. A total of 478 species clusters contains at least 2 genomes.

**Figure S3.**
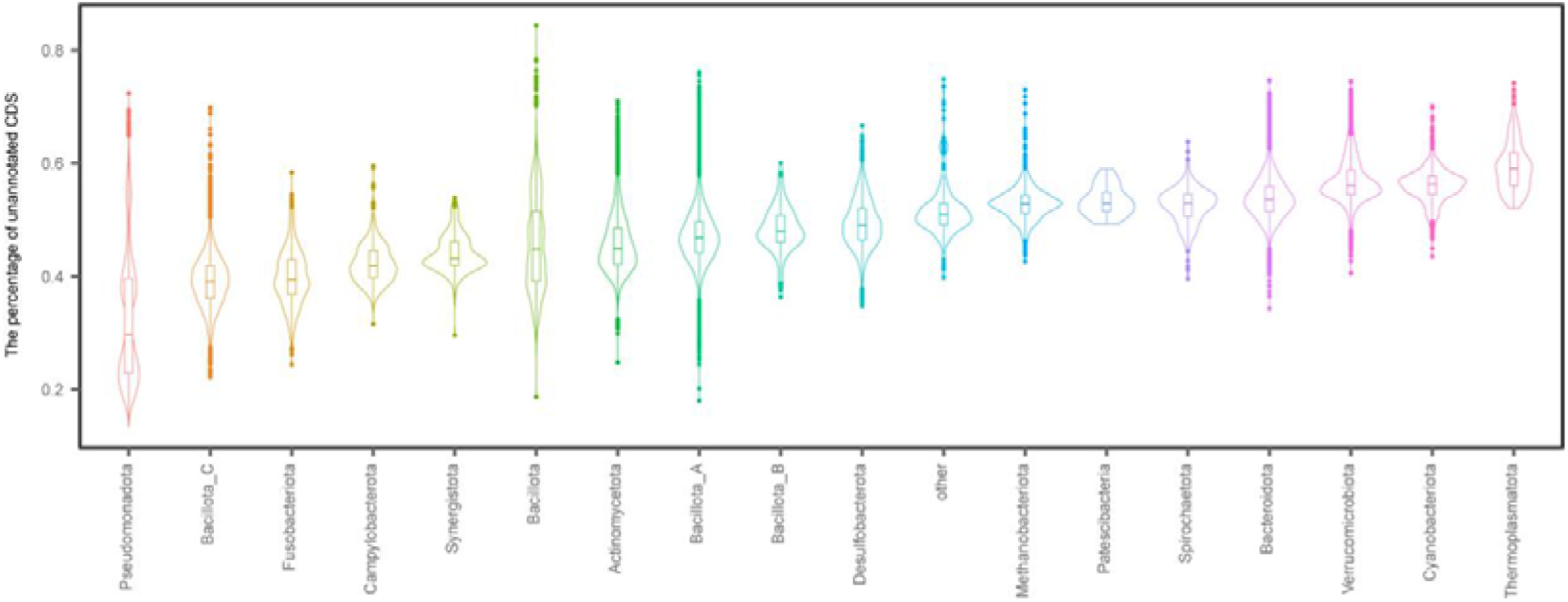
Phylum-level proportion of functionally annotated versus unannotated CDSs. The bar plot shows the percentage of CDS with unassigned function within each phylum. The phyla comprising < 10 of each genome set was collapsed into other. The boxplots display the median values (center lines), interquartile ranges (IQRs, represented by the box edges), and whiskers, which extend to the most extreme data points within 1.5 times the IQR from the lower and upper quartiles.

**Figure S4.**
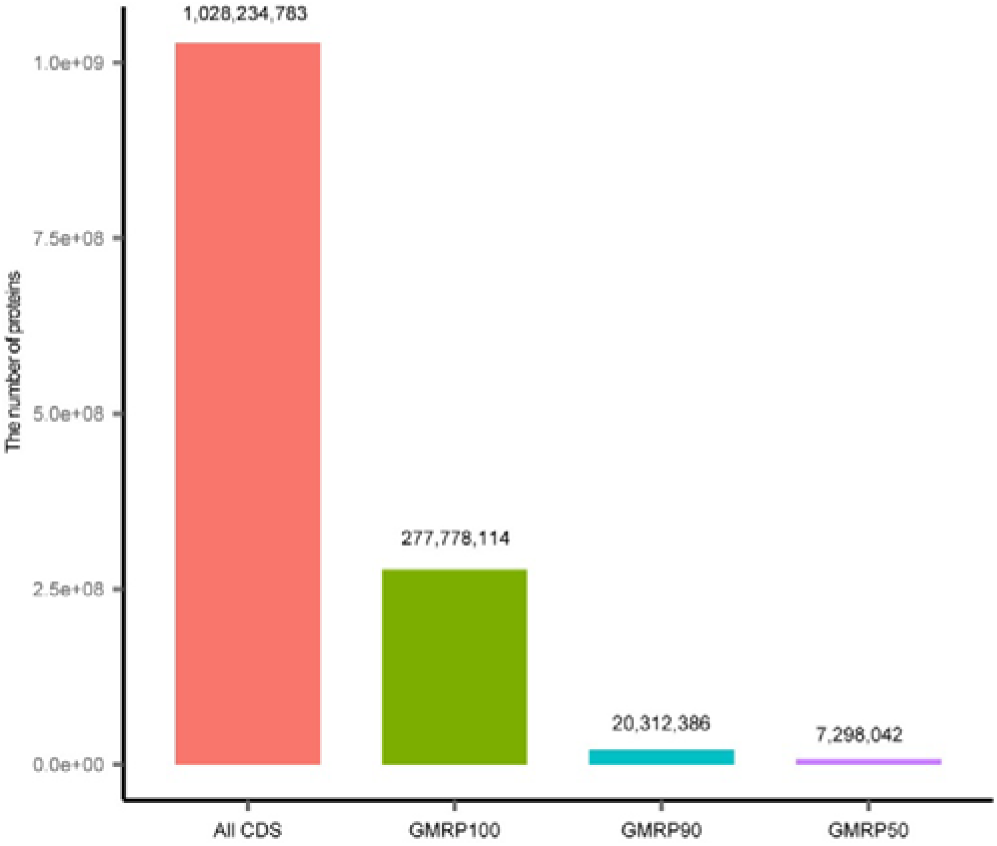
Clustering of GMR protein functions. The representation of protein classes characterized by GMR after clustering at amino acid similarity thresholds of 100%, 90%, and 50%, respectively.

**Figure S5.**
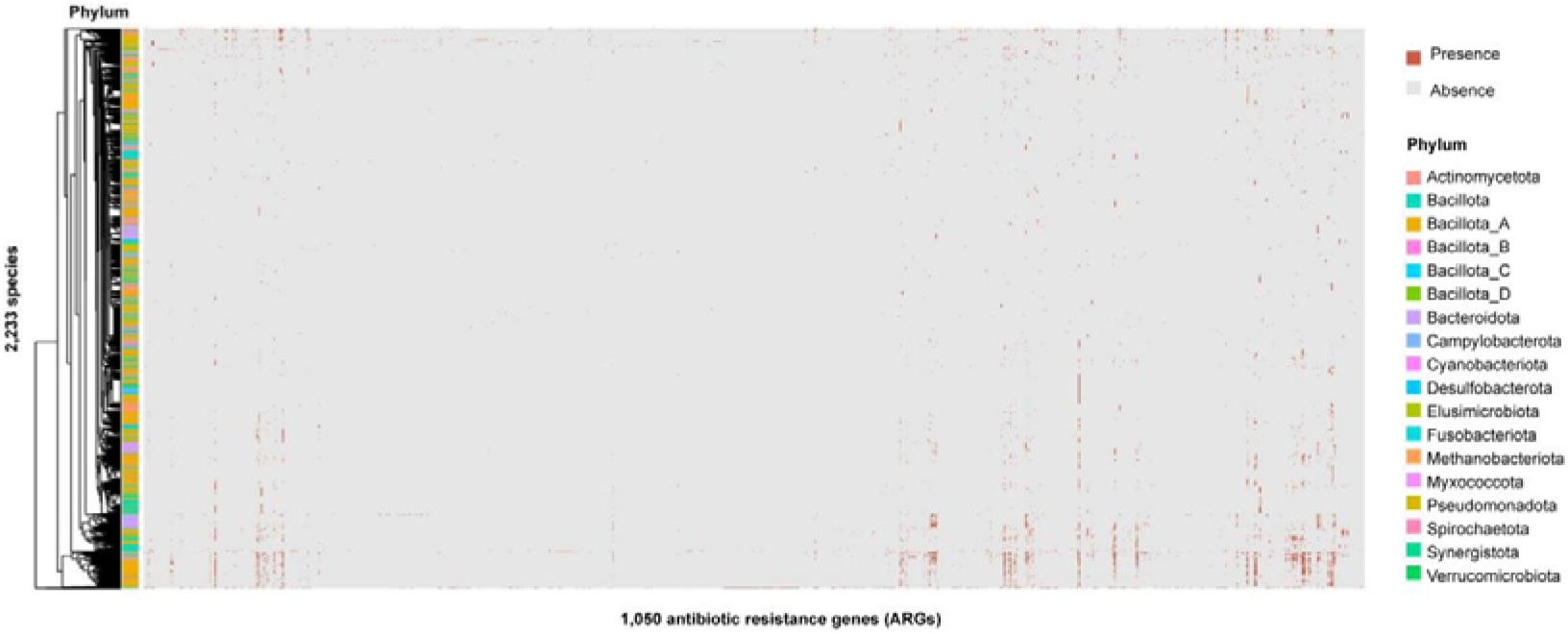
Detection of antibiotic resistance genes (ARGs) in GMR-associated microorganisms. A total of 1,050 ARGs were identified across 2,233 species. Rows represent species, columns represent genes, and phylum classifications are indicated by corresponding colors.

**Figure S6.**
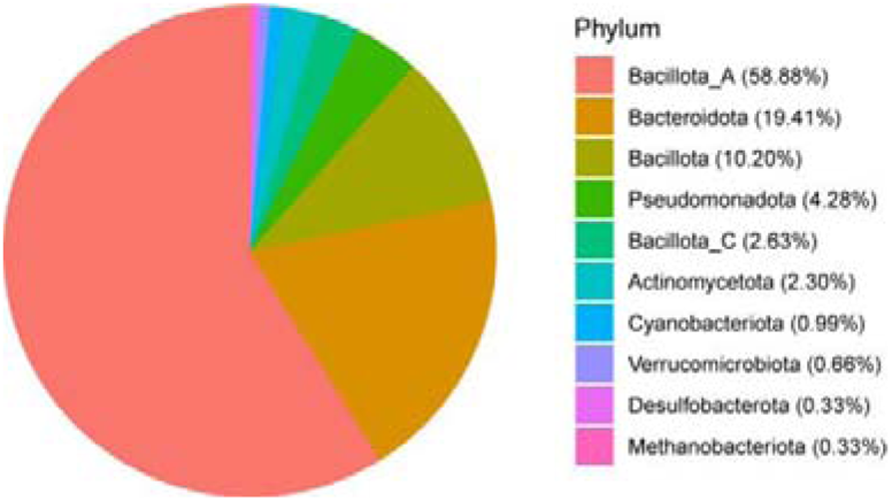
Phylum-level classification of 304 predominant species. Bacillota_A dominate the composition (58.88% of total species).

**Figure S7.**
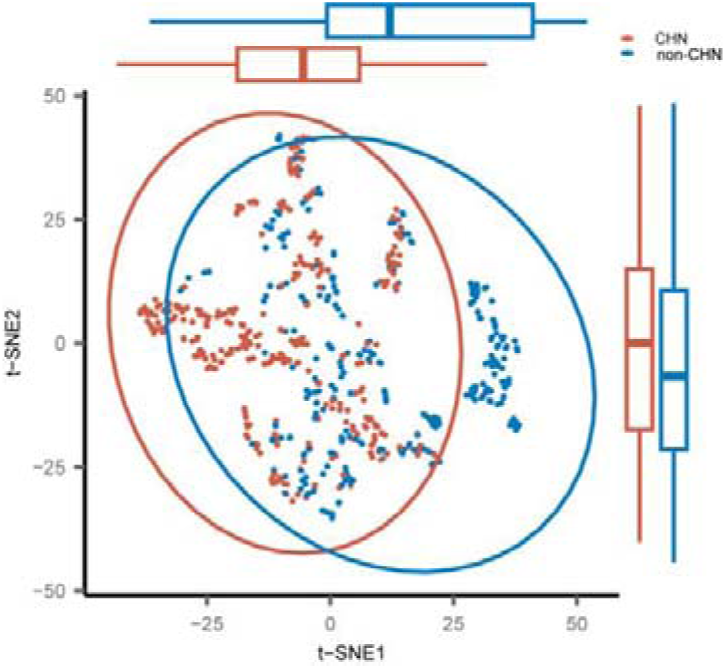
Population-level genetic structure of *Prevotella copri*. The scatter plot represents tSNE coordinates, with corresponding boxplots showing distribution patterns along each tSNE axis.

**Figure S8.**
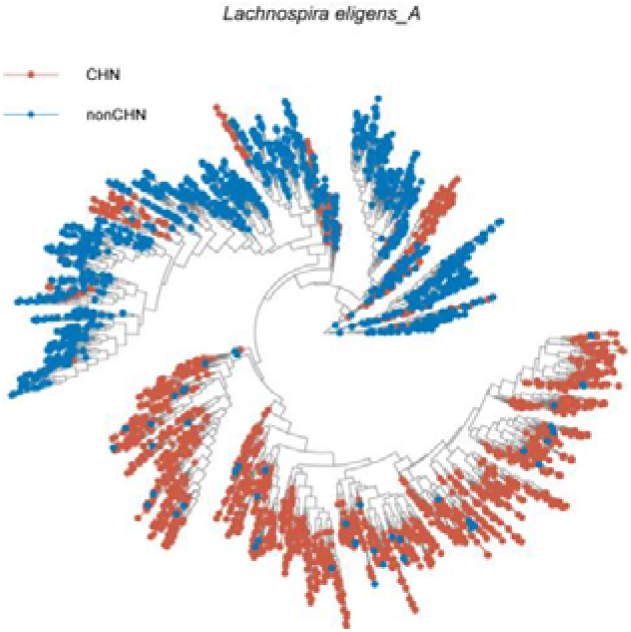
Phylogenetic tree of *Lachnospira eligens_A* based on gene presence/absence patterns. MAGs from Chinese individuals form a distinct cluster.

**Figure S9.**
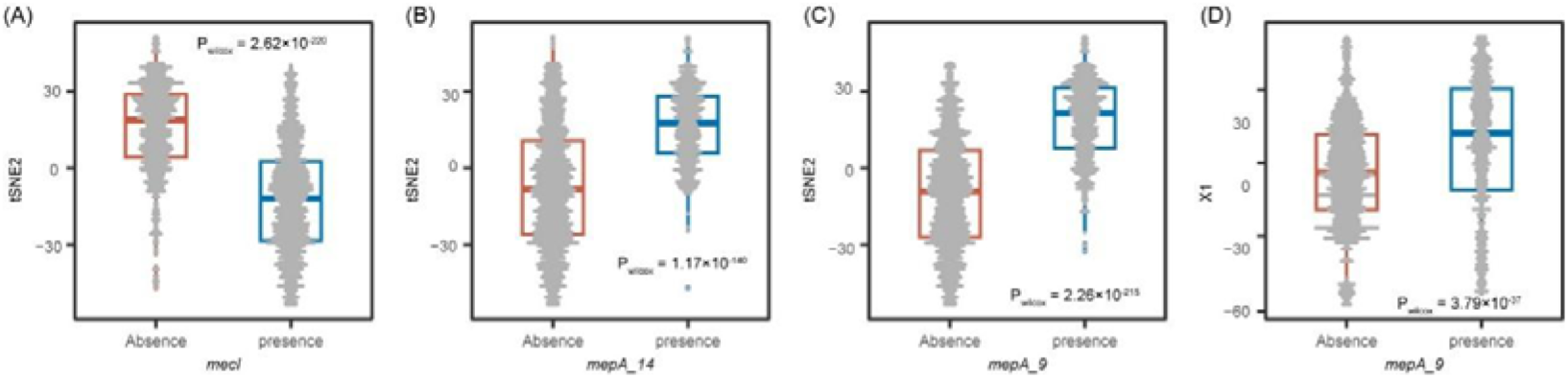
Association between gene presence/absence and genetic structure in *Lachnospira eligens_A*. Panels A–D depict correlations of *mecl*, *mepA_14 and mepA_9* with tSNE coordinates, respectively. Red represents absence and blue represents presence. The boxplots show the median, first quartile (25th percentile), and third quartile (75th percentile), with whiskers extending to the most extreme data points within 1.5 times the interquartile range. Each dot represents a MAG derived from an individual.

**Figure S10.**
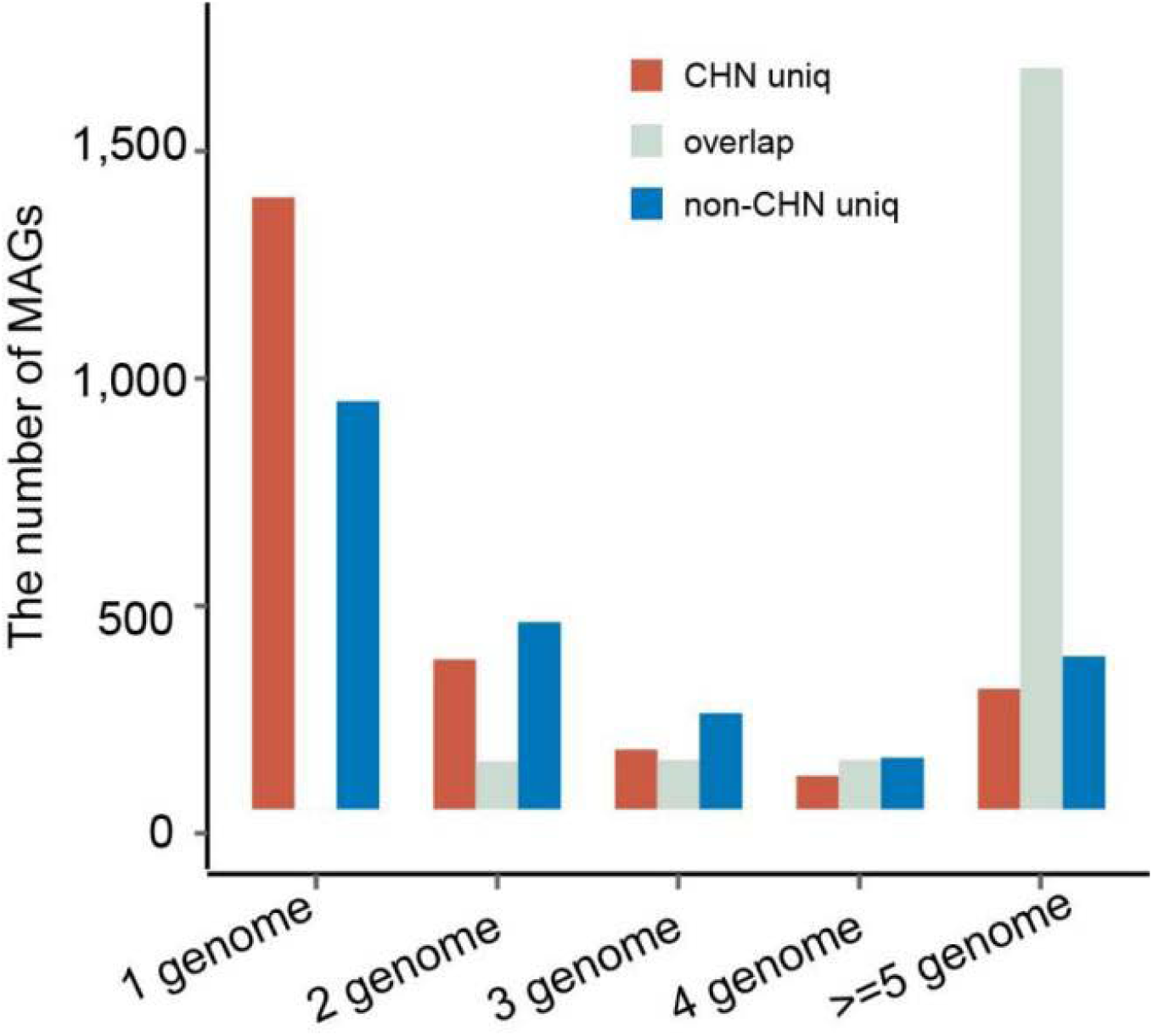
Distribution of microbial genomes across species-level clusters. The bar plot shows the number of genomes unique to Chinese and non-Chinese populations in each species cluster.

**Figure S11.**
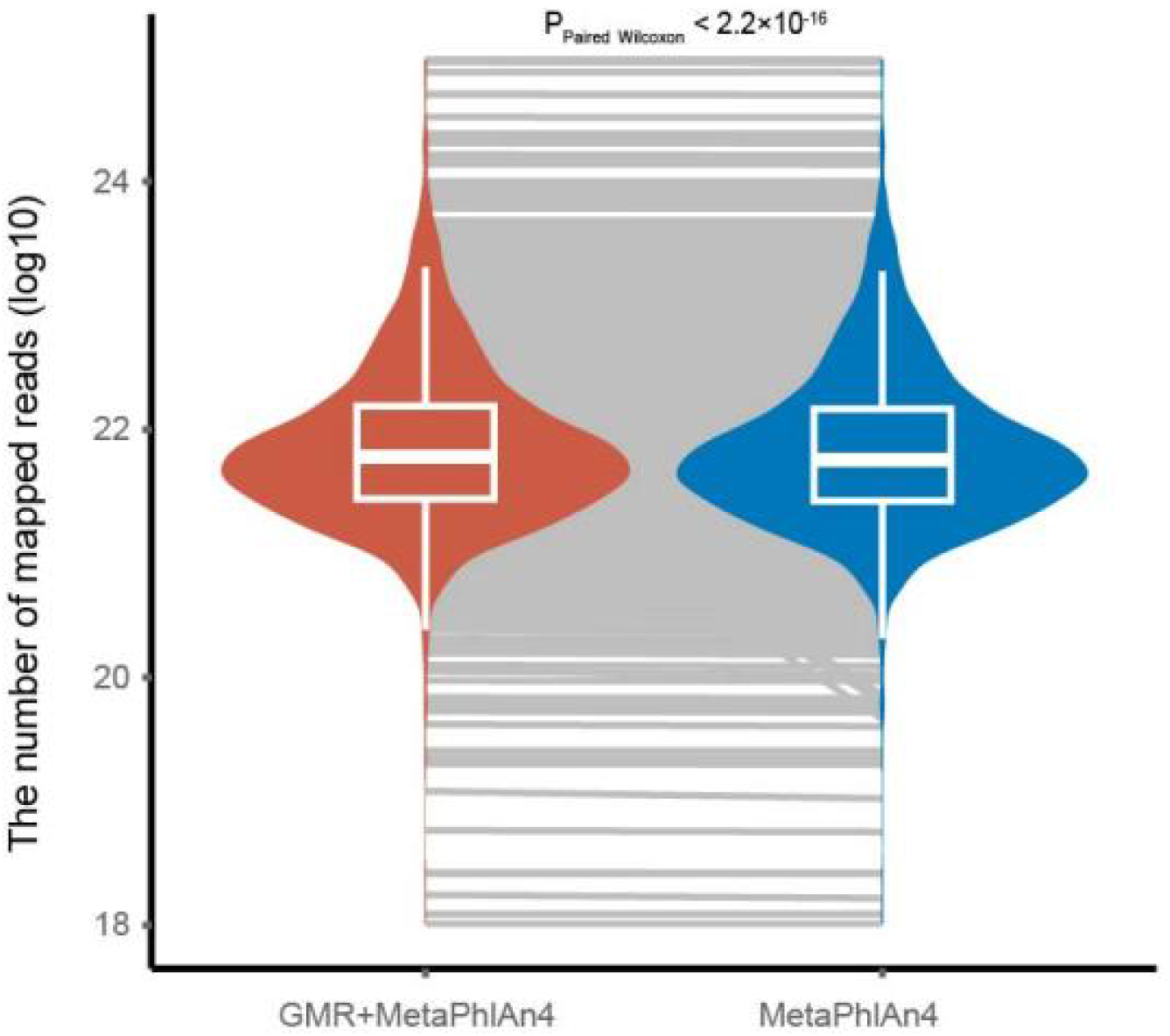
Enhancement of the reads mapping rate for an independent Chinese-derived dataset using the new database. After integrating the GMR into the MetaPhlAn4 database for species identification, an improvement in the reads mapping rate was observed (P_Paired wilcoxon_ < 2.2×10^-16^).

**Figure S12.**
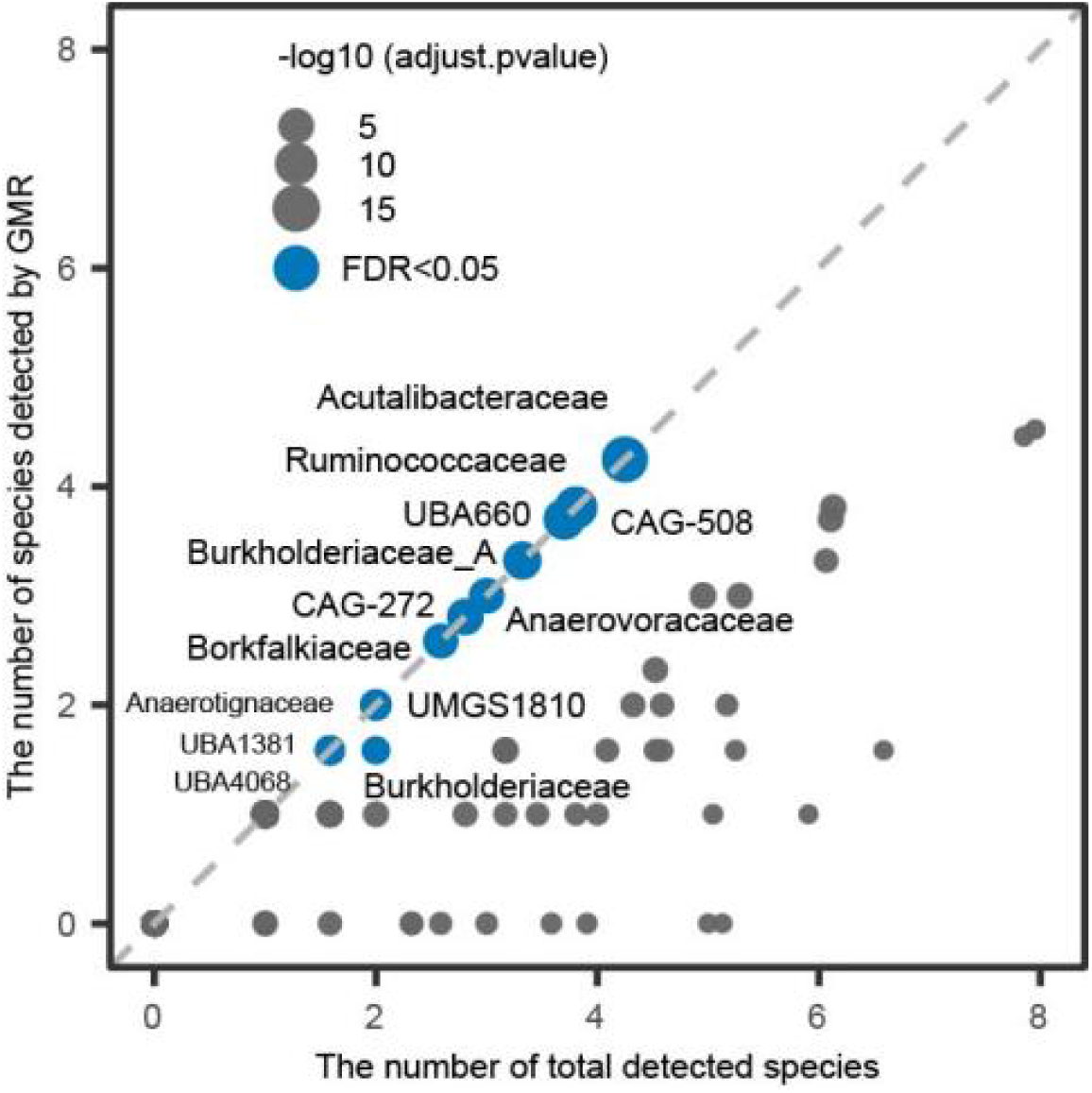
Summary of microbial species showing significant improvement at the family level using the new database. A total of 326 novel species were detected after integrating the GMR into the MetaPhlAn4 database. The X-axis represents the total number of species detected within each family, while the Y-axis shows the number of newly identified species. Each dot represents a family, with the size of the dot indicating the FDR value from the hypergeometric test. Blue dots represent families with an FDR < 0.05. Among the 92 families containing newly detected species, 13 showed a significant improvement at an FDR threshold below 0.05, and these families are labeled accordingly.

**Figure S13.**
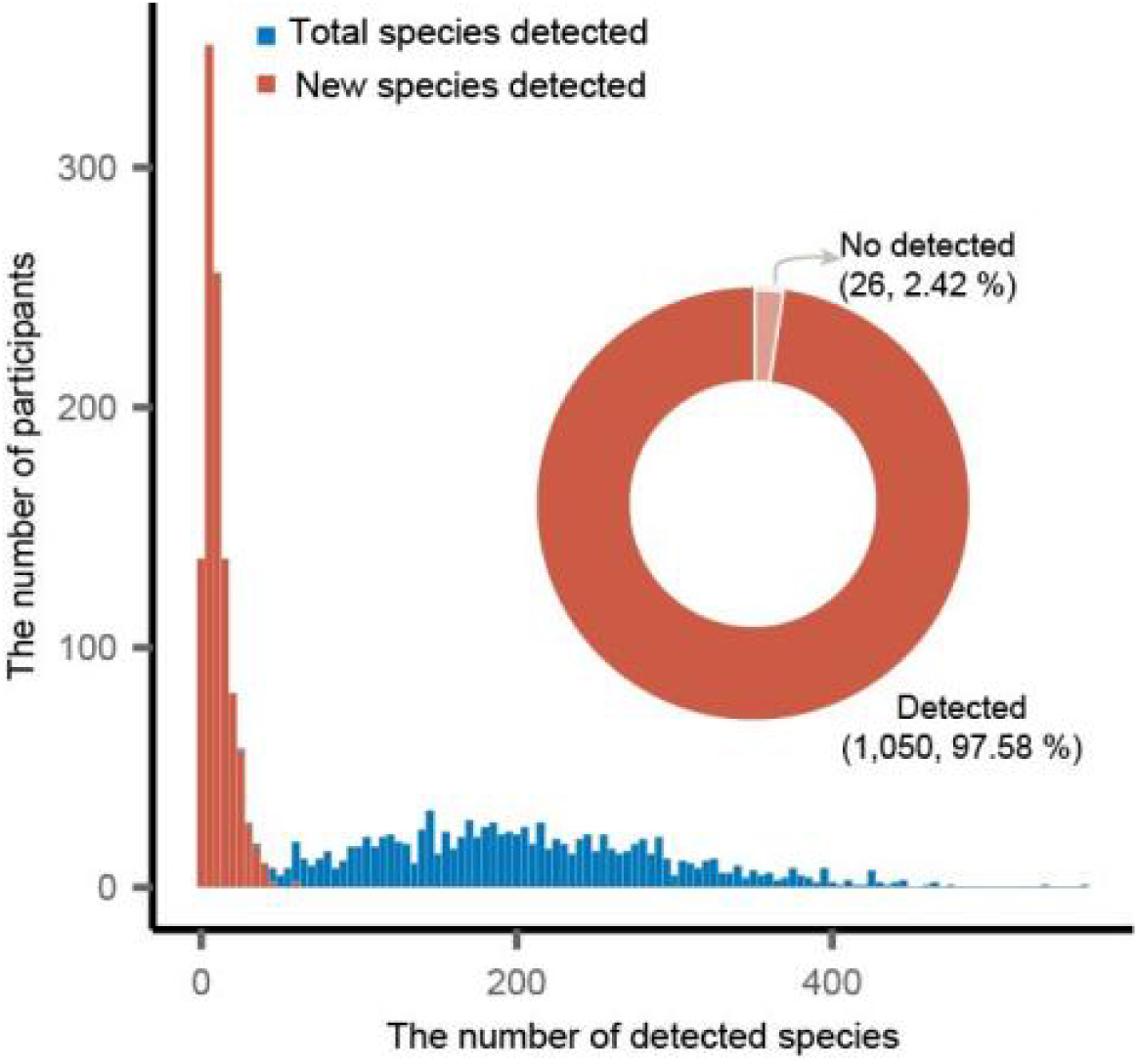
Summary of microbial species profiled in a cohort of 1,076 participants. The blue bars represent the total number of detected species, while the red bars indicate the number of newly classified species in the GMR. On average, 205 species were detected per sample, with 11 belonging to newly classified species in the GMR. The pie chart on the right illustrates the proportion of samples with at least one newly detected species, showing that 97.58% of the samples contained at least one new species.

**Figure S14.**
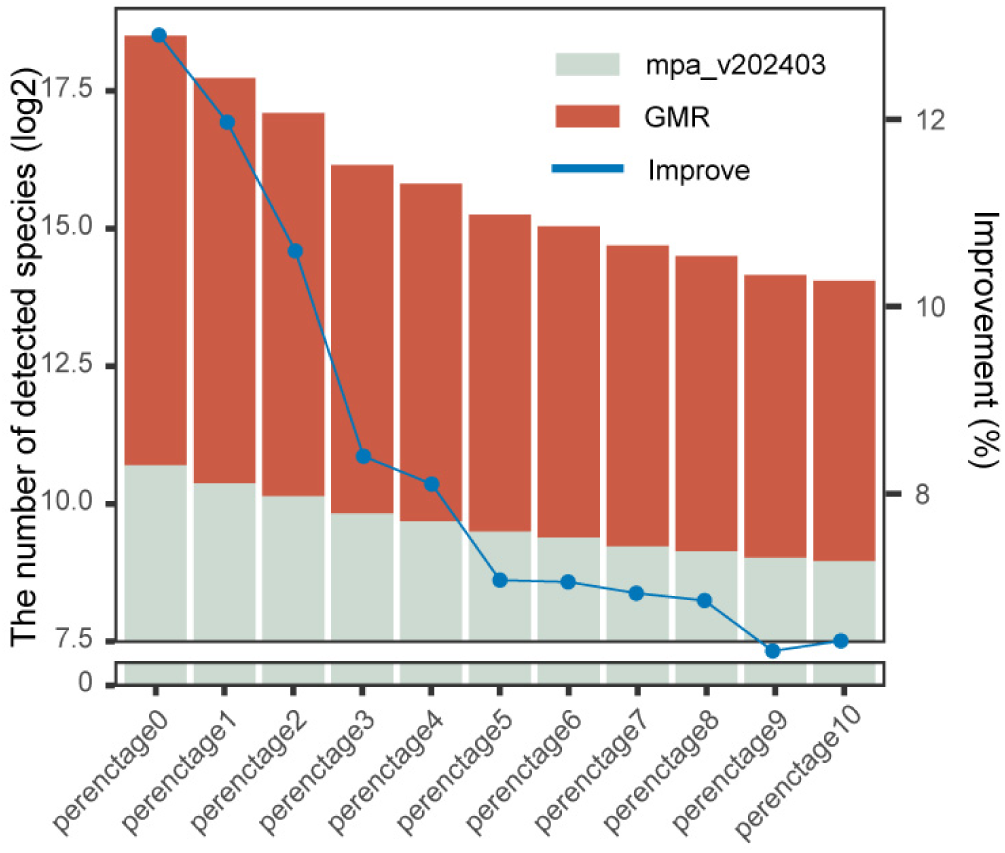
Taxonomic profiling at the species level using the newly constructed database in a German cohort with diverse filtering cutoffs. A total of 149 fecal metagenomic samples were analyzed. The X-axis shows different filtering cutoffs, from 0% to 10%. The left Y-axis (bar chart) represents the number of detected species, with the red portion highlighting the contribution of newly identified species in the GMR. The right Y-axis (blue line) represents the improvement in species detection achieved by including the newly identified species from the GMR.

**Figure S15.**
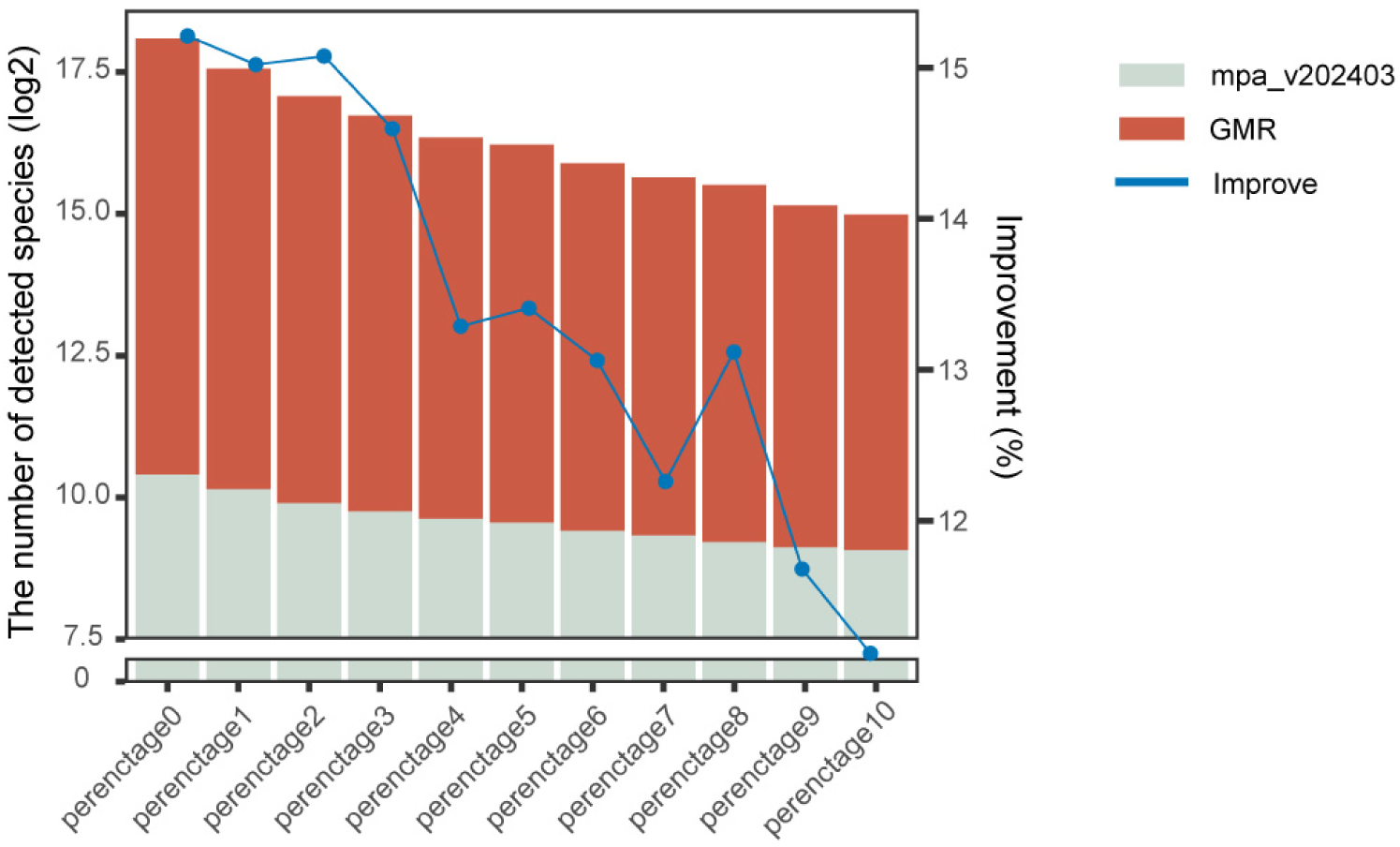
Taxonomic profiling at the species level using the newly constructed database in a Congolese cohort with diverse filtering cutoffs. A total of 179 fecal metagenomic samples were analyzed. The X-axis shows different filtering cutoffs, from 0% to 10%. The left Y-axis (bar chart) represents the number of detected species, with the red portion highlighting the contribution of newly identified species in the GMR. The right Y-axis (blue line) represents the improvement in species detection achieved by including the newly identified species from the GMR.

**Figure S16.**
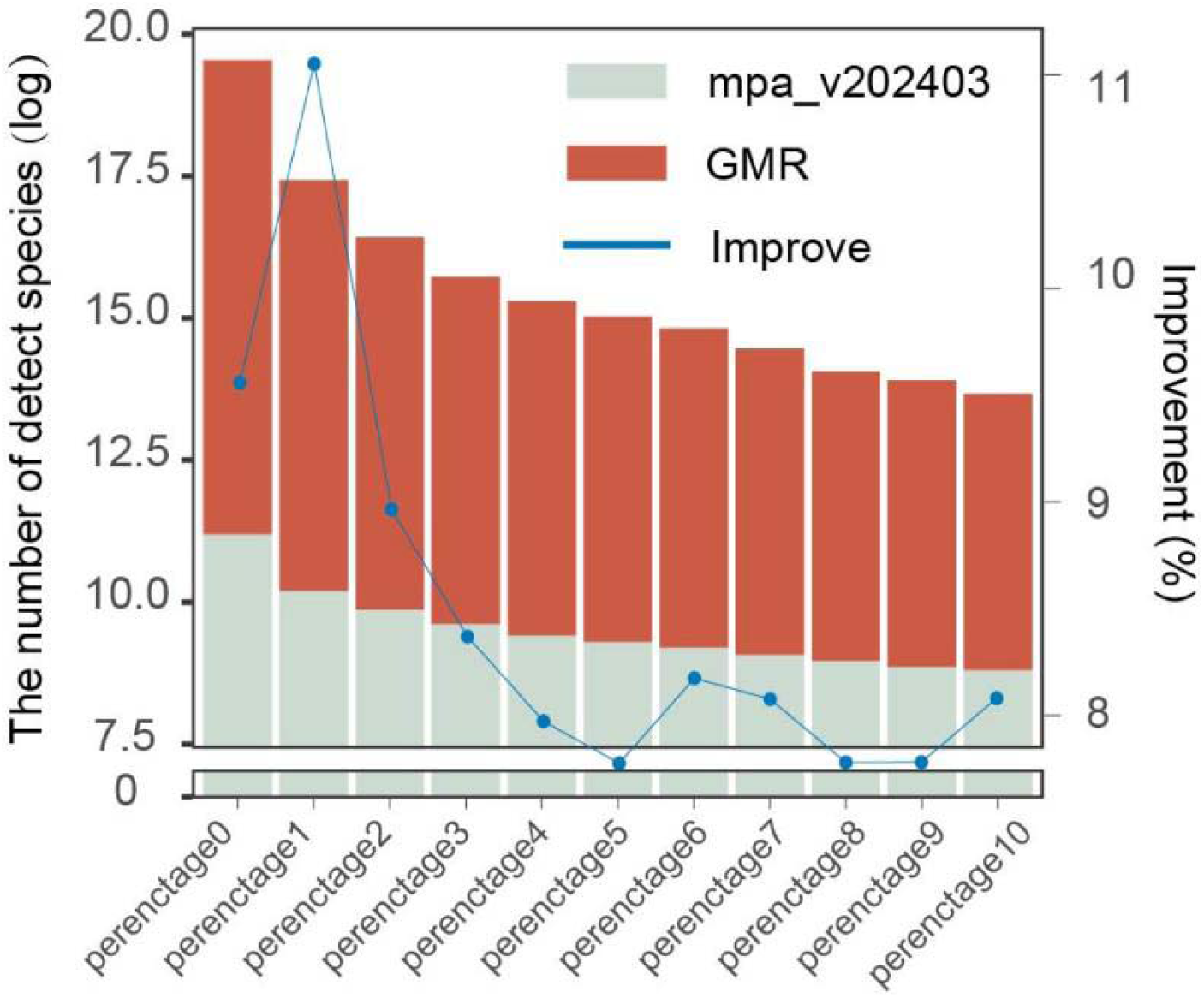
Taxonomic profiling at the species level using the newly constructed database in the CGMR cohort with diverse filtering cutoffs. A total of 3,234 fecal metagenomic samples were analyzed. The X-axis shows different filtering cutoffs, from 0% to 10%. The left Y-axis (bar chart) represents the number of detected species, with the red portion highlighting the contribution of newly identified species in the GMR. The right Y-axis (blue line) represents the improvement in species detection achieved by including the newly identified species from the GMR.

**Figure S17.**
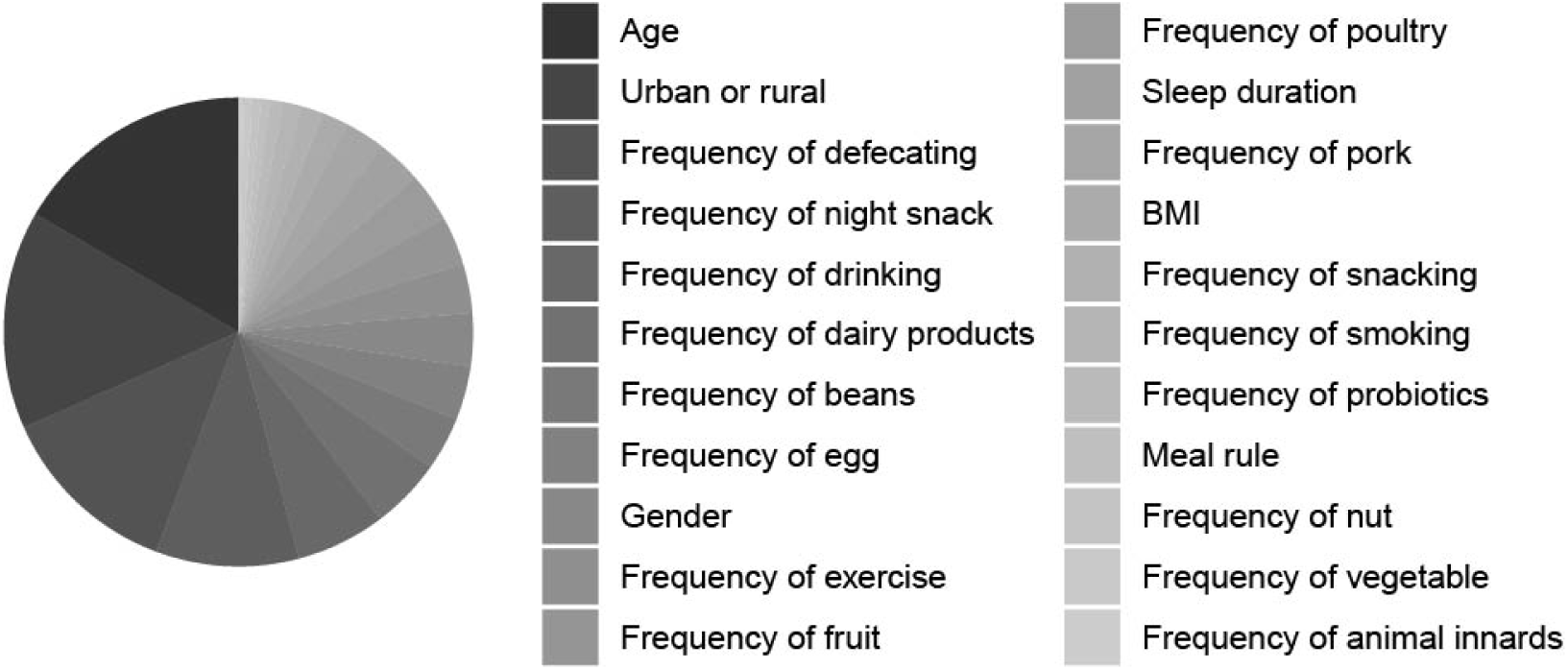
Summary of host phenotypic associations with the abundance of newly profiled species. Pie chart illustrates the percentage of phenotypes that each bacterial species is significantly associated with. Different colors represent different phenotypes.

**Figure S18.**
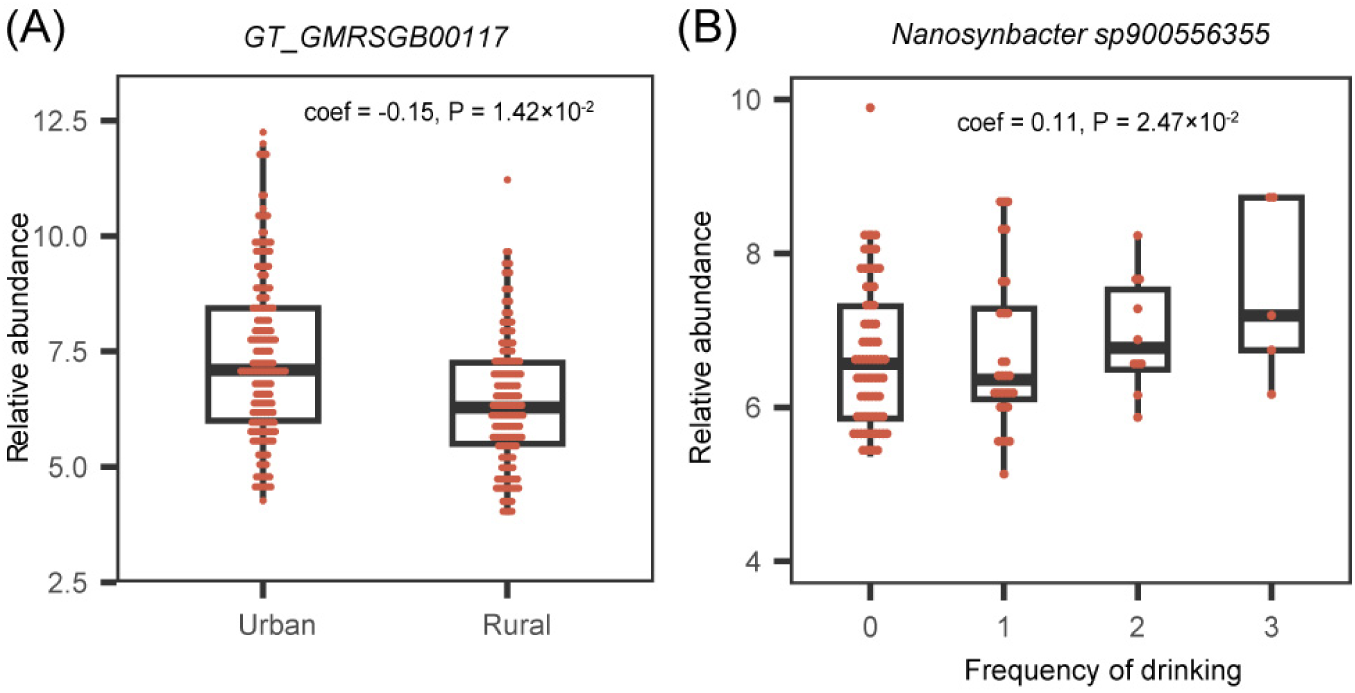
Associations between species abundance and phenotypes in the CGMR cohort. (**A**). The relative abundance of *GT_GMRSGB00117* is observed to be lower in rural areas**. (B).** The relative abundance of *Nanosynbacter sp900556355* iincreases with increasing drinking frequency. The box plot compares species abundance across different defecation frequencies. The boxes represent the median, first, and third quartiles (25th and 75th percentiles), with whiskers extending to the most extreme data points within 1.5 times the interquartile range. Each dot represents a participant.

## Table Legends

**Table S1.** The summary of assembly statistics of the 478,588 human gut microbial genomes used to generate the Global Microbiome Reference (GMR) collection.

**Table S2.** The metadata of 6,664 representative species of GMR and their novelty status.

**Table S3.** Comparison of the number of genomes for shared species between Chinese and non-Chinese populations.

**Table S4.** The summary of CDS identified per genome in the GMR and the proportion of CDS with unknown function.

**Table S5.** COG functional annotation of GMRP90.

**Table S6.** Number of antibiotic resistance genes (ARGs) detected per species.

**Table S7.** Gene distribution on MAGs with the highest number of detected antibiotic resistance genes (ARGs).

**Table S8.** Distribution of antibiotic resistance genes (ARGs) detected in Chinese and non-Chinese populations.

**Table S9.** Number of genomes per species in the GMR database.

**Table S10.** Comparison of genetic structure between populations.

**Table S11.** Enrichment of species with differences in genetic structure between populations.

**Table S12.** Number of the core CDS and marker genes identified for species with genome greater than or equal to 3.

**Table S13.** Species information ultimately incorporated into the MetaPhlAn4 database.

**Table S14.** Families with the highest species enrichment detected after GMR Integration.

**Table S15.** The summary of species abundance profiling of the CGMR database using customized database.

**Table S16.** Sum of abundances of unique species detected in each sample by CGMR.

**Table S17.** The association between the abundance of novel species detected using our personalized database and phenotypes of CGMR(fdr<0.05).

**Table S18.** The correlations of species abundance and phenotype found were validated in another independent cohort.

