## Supplementary figures and images for "Expanded gut microbial genomes from Chinese populations reveal population-specific genomic features related to human physiological traits"

### FigureS1

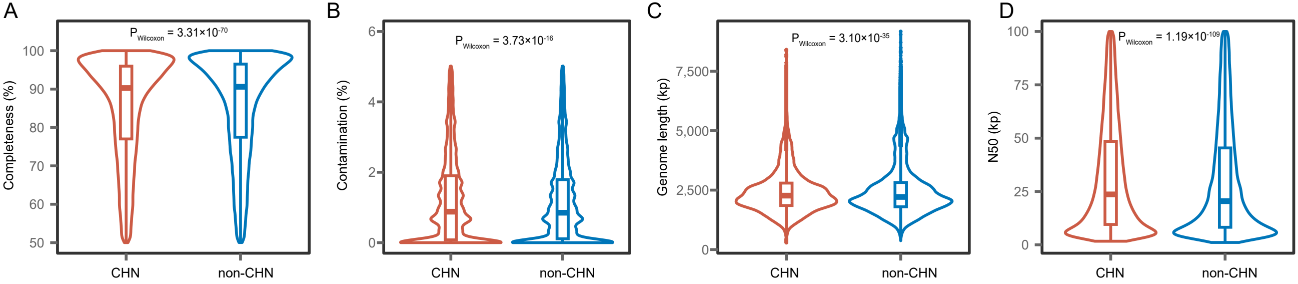

### FigureS2

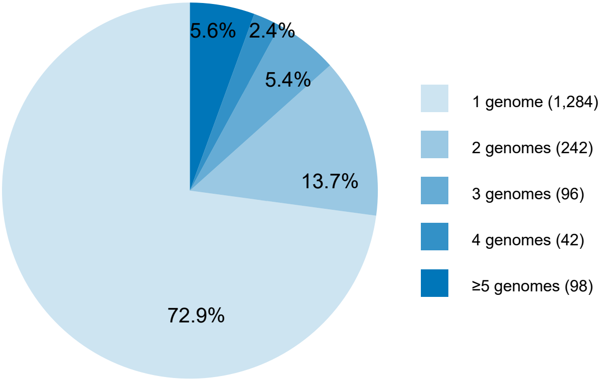

### FigureS3

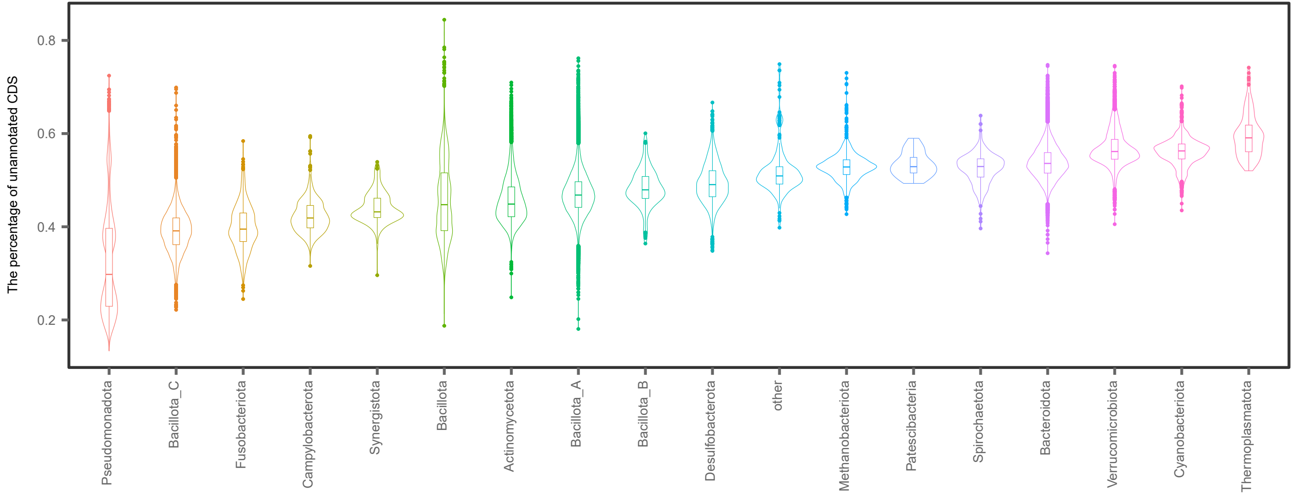

### FigureS4

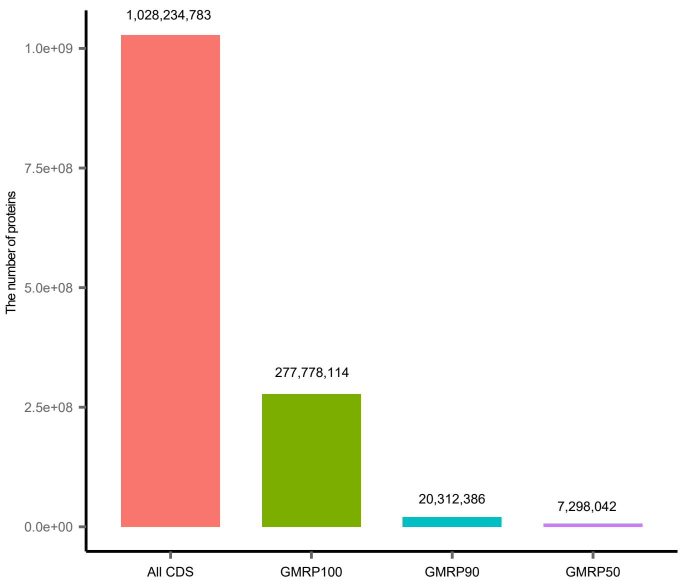

### FigureS5

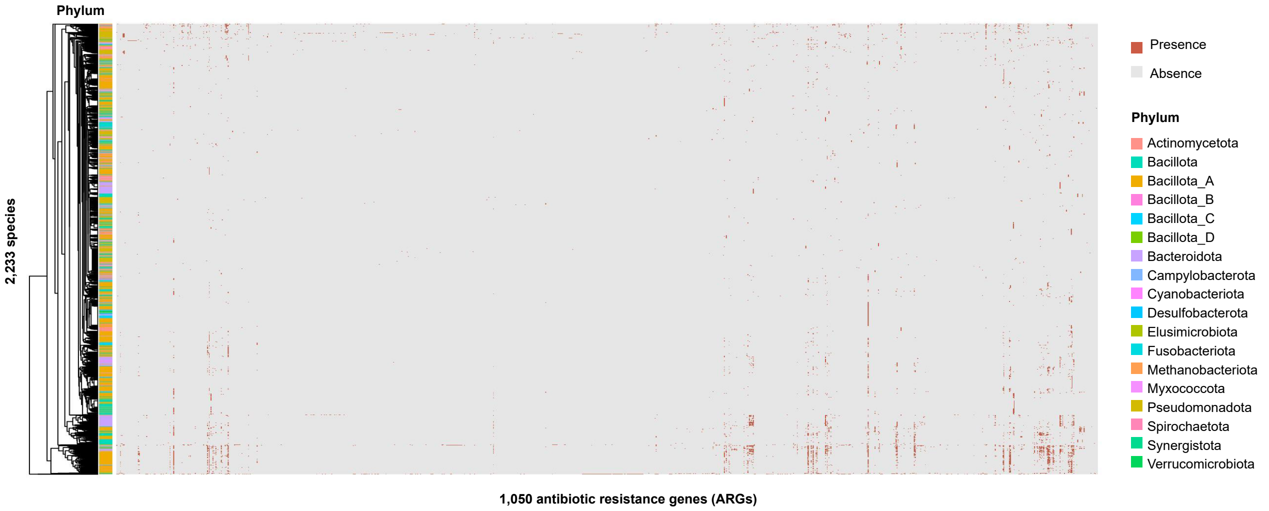

### FigureS6

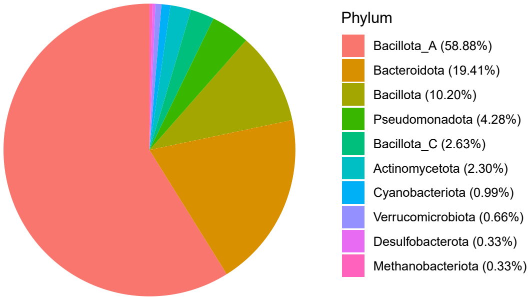

### FigureS7

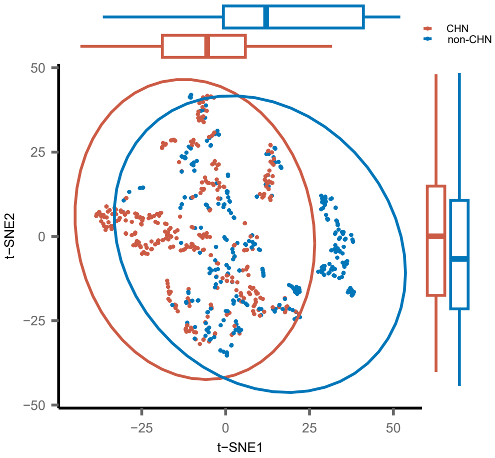

### FigureS8

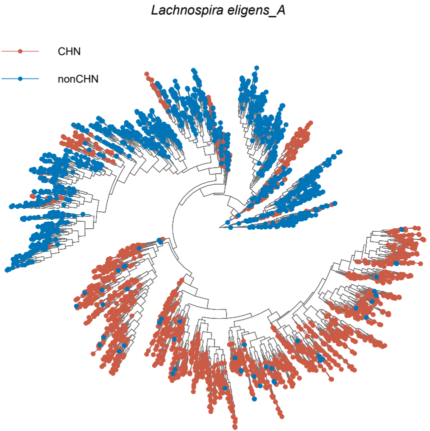

### FigureS9

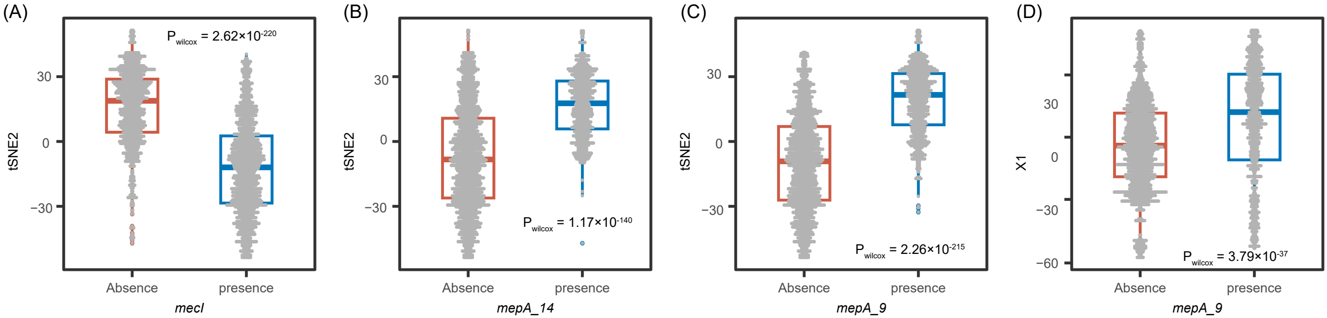

### FigureS10

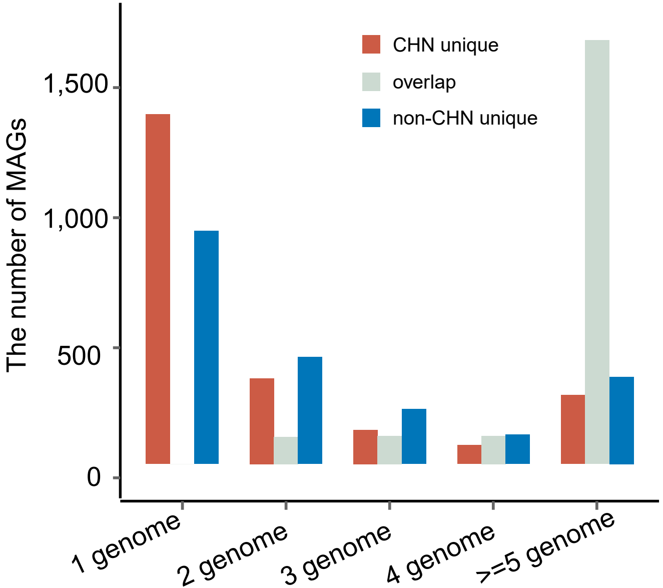

### FigureS11

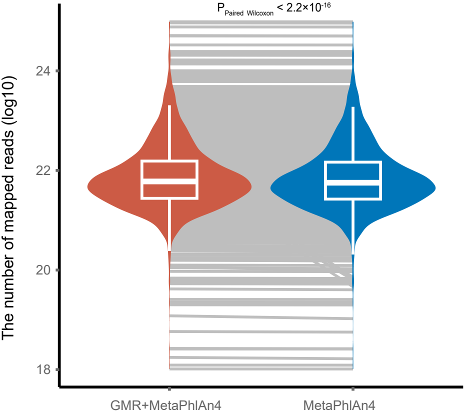

### FigureS12

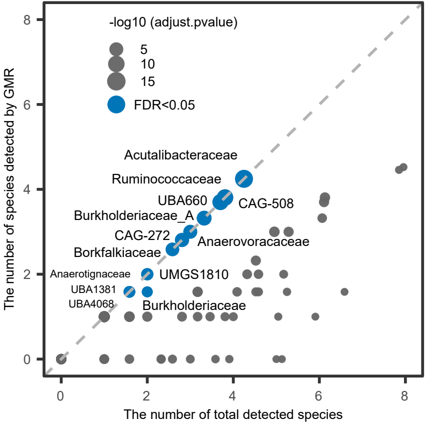

### FigureS13

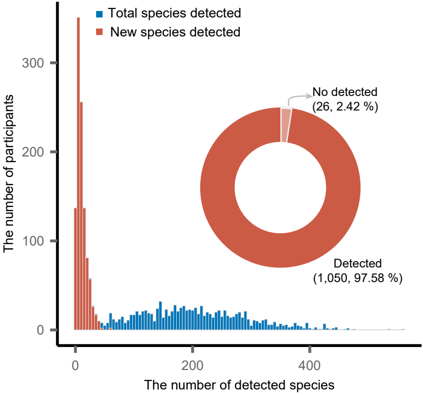

### FigureS14

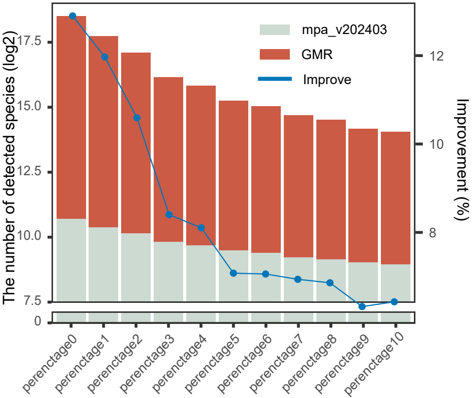

### FigureS15

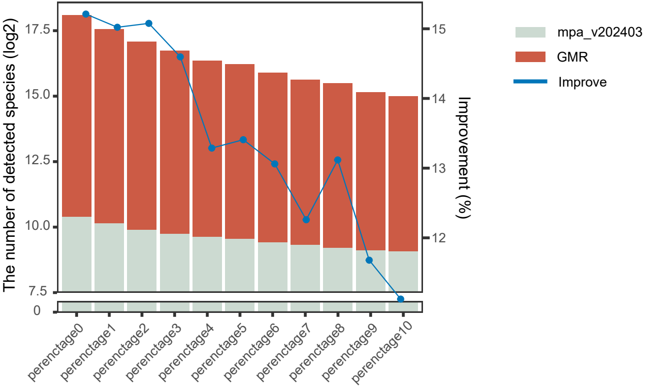

### FigureS16

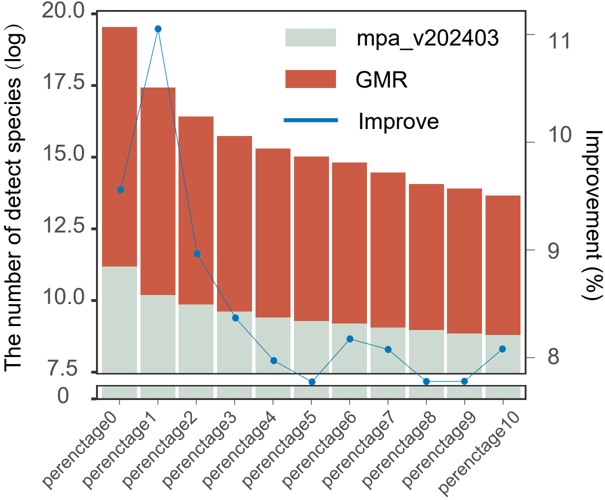

### FigureS17

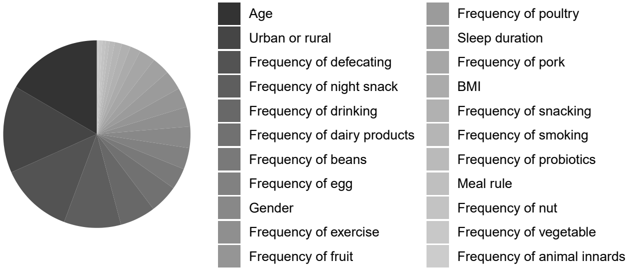

### FigureS18

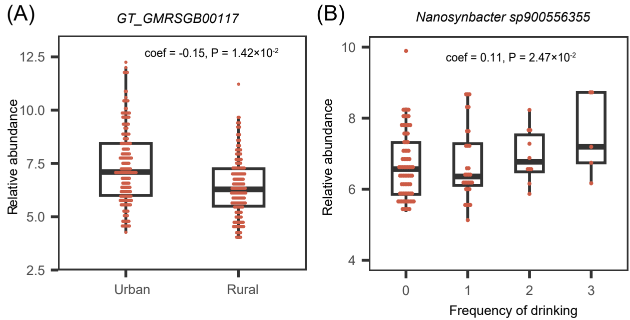
